# An AAV model of tauopathy targeting the mouse dentate gyrus reveals greater trans-synaptic tau spreading with aggregation-prone tau variants

**DOI:** 10.64898/2026.11.09.762480

**Authors:** Pauline Léal, Charlotte Duwat, Cécile Adamski, Marie-Claude Gaillard, Gwennaëlle Auregan, Martine Guillermier, Sueva Bernier, Noëlle Dufour, Fanny Petit, Caroline Jan, Anne-Sophie Hérard, Déborah Fourmy, Mehdi Kabani, Gilles Bonvento, Karine Cambon, Alexis-Pierre Bemelmans

## Abstract

Alzheimer’s disease is characterized by progressive memory and cognitive decline, which is strongly correlated with the anatomical progression of tau pathology. Resulting from the intraneuronal aggregation of misfolded tau protein, tauopathy propagates in an anterograde manner between synaptically connected neurons through a prion-like mechanism. While glial cells, particularly microglia, are implicated in this process, the precise mechanisms underlying tauopathy propagation remain poorly understood. A critical unresolved question is whether aggregated or soluble pathologic tau forms mediate this spread. Here, we show *in vivo* that tau spreading scales with the aggregation propensity of the tau variant expressed. We developed a mouse model using adeno-associated virus (AAV)-mediated gene transfer to overexpress distinct tau forms in the dentate gyrus, alongside a reporter gene. This design enabled discrimination between transduced neurons and those secondarily affected by pathology propagation. Leveraging contralateral neuronal projections, we quantified propagation levels. To investigate the role of microglia and of extracellular vesicles, we analyzed microglial morphology in response to different tau forms and measured tau protein levels in extracellular vesicles isolated from injected mouse brains. We first confirmed the presence of vector-derived tau protein in non-transduced neurons, validating our model for studying tau propagation. We then found that aggregation-prone tau variants were associated with a greater number of tau-positive neurons in the contralateral hippocampus than wild-type tau. Additionally, extracellular vesicles from brains injected with aggregated tau contained higher levels of tau protein. Given that insoluble tau aggregates represent the ultimate stage of tauopathy, our findings suggest that propagation occurs at advanced stages of tau pathology, and that extracellular vesicles may contribute to the transfer of pathological tau, although the cellular origin of these vesicles was not determined in this study. Furthermore, our model provides a robust experimental paradigm for exploring therapeutic strategies aimed at inhibiting tauopathy propagation.

## INTRODUCTION

Although Alzheimer’s disease (AD) is consistently marked by the coexistence of two proteinopathies, amyloidosis and tauopathy, cognitive impairment correlates more closely with the burden and anatomical distribution of tau pathology than with amyloid deposition.^1^ The first evidence supporting the propagation of tau protein between anatomically connected neurons in AD was provided by the seminal histopathological studies of Braak and colleagues. Their staging system, based on post-mortem brain analyses, revealed a stereotypical and progressive pattern of neurofibrillary tangle spread that followed neuroanatomical connectivity, suggesting transneuronal transmission of pathological tau.^2^ Notably, this progression of tau pathology correlates strongly with the emergence and severity of cognitive symptoms, more than amyloid deposition.^1,3^ Tau PET imaging subsequently showed, in living patients, a spatiotemporal pattern of tau accumulation that recapitulates the Braak stages, and that the anatomical connectivity of a region predicts where tau appears next.^4,5^ Direct evidence for transneuronal transfer, however, comes from *in vitro* and *in vivo* experimental models.^6–9^ These findings have provided critical support for the concept of tau as a propagating proteinopathy and a key driver of cognitive decline in AD. Thus, slowing or stopping the spreading of tauopathy would be an essential leverage to set-up a disease modifying treatment able to influence the progression of cognitive symptoms. Deciphering the molecular mechanisms of tau spreading is essential, first to understand the complex dynamics of this neurodegenerative disease, then to develop innovative and targeted strategies of treatment. Tau propagation has been first described as a phenomenon following a prion-like spreading leading to the misfolding of endogenous tau after being in contact with pathogenic misfolded tau.^10^ This prion-like propagation has been shown both in *in vitro* models^6^ and in some *in vivo* models.^7^ Transneuronal spreading is described as the spreading of tau from the presynaptic to the postsynaptic neuron. The ability of tau to spread from a neuron to another at the synapse has been demonstrated *in vitro* and *in vivo.*^6,8,9^ Some studies also showed that tau was able to spread from neuron to neuron after injection of lentiviral vectors expressing either hTau^WT^ (human tau wild-type) or hTau^P301L^ (human tau with P301L mutation) in the hippocampus of rats^11^ or injection of AAV vectors expressing tau in the entorhinal cortex of mice.^12,13^

Tau spreading mechanisms have been thoroughly studied and several mechanisms involved in the secretion of tau by presynaptic neurons and in the internalization of tau by postsynaptic neurons have been identified. Once secreted, tau can be taken up by postsynaptic neurons through several mechanisms. One of the most characterized to date is macropinocytosis mediated by heparan sulfate proteoglycans (HSPGs), which facilitates nonspecific endocytosis of tau, particularly in aggregated form.^14–16^ Furthermore, the LRP1 receptor (Low-Density Lipoprotein receptor-related Protein 1) is involved in the uptake of monomeric tau and its inhibition significantly reduces the spreading of pathology in experimental models.^17,18^ Other entry mechanisms have been proposed, including clathrin-dependent endocytosis, endocytosis mediated by other receptors such as muscarinic receptors, as well as internalization *via* transient membrane structures induced by tau aggregation.^18,19^ Other studies also suggest that pathological tau may directly transfer to recipient neuron through tunneling-nanotubes.^20–22^ Furthermore, tau has been found in different types of extracellular vesicles such as exosomes and ectosomes.^23,24^ Tau can be found free in the extracellular space and it is now well established that neuronal activity promotes the secretion of tau from presynaptic terminals. However, it has been suggested that tau associated with extracellular vesicles spreads more efficiently than free tau.^25^ Although no predominant mechanism has yet been identified in pathological condition, synaptic contacts have now been shown to facilitate this propagation, likely by increasing the efficiency of tau transfer from one neuron to another.^26^ This release is dependent on synaptic activity and activation of glutamatergic receptors, as well as on overall increase in neuronal activity, which induces the exocytosis of soluble form of tau, even in the absence of neurodegeneration.^27–30^

Interestingly, several lines of evidence involve microglia, the main immune cell-type of the central nervous system. First, several gene variants associated with increased risk for AD are exclusively expressed by microglia within the central nervous system.^31–33^ Beyond their function of phagocytosing Aß and tau aggregates, microglia may also actively contribute to the dissemination of tau pathology.

First, Bolòs *et al.* showed that microglia are able to internalize tau protein both *in vitro* and *in vivo*.^34^ Another study showed the ability of microglia to phagocytose extracellular tau and partially degrade it without preventing its seeding property.^35^ Using an AAV-based model of tauopathy, Asai *et al.* demonstrated that depletion of microglia *in vivo* and inhibition of exosomes biogenesis significantly reduce tau spreading in mouse models of AD.^12^ A link has also been made between microglia, amyloidosis and spreading of tau. Clayton *et al.* showed that amyloidosis and microglia have a synergic effect in worsening the spreading of tau *via* microglia activation in presence of amyloidosis which increases exosomes release and tau spreading.^36^ Finally, transcriptomic analyses have identified specific microglial subpopulations associated with tau pathology, suggesting functional specialization in the propagation of tauopathy.^37^ These data highlight the active role of microglia in the propagation of tauopathy and suggest that targeting microglial mechanisms could be a promising therapeutic strategy to slow the progression of AD.

The ability of tau to spread from one neuron to another involves a lot of different factors. Among those, the propensity of tau to spread regarding its degree of aggregation has not really been assessed yet. To tackle this question, we set up innovative models of tauopathy specifically designed to study the spreading of the pathology. In a previous work, we showed that the use of AAVs containing both a GFP reporter gene and a V5 tag inserted in tau’s open reading frame, separated by a P2A sequence allowing the cleavage of both peptides during translation, is a valuable tool to distinguish AAV-transduced neurons from the ones in which tauopathy occurred by spreading.^38^ We injected this AAV in the vitreous of the mouse eye to target the retinal ganglion cells (RGCs) and observed a good transduction of RGCs and the spreading of tauopathy in the brain. However, in this experimental paradigm, the spreading remained very limited, appearing only in old animals, and suggesting that there are intrinsic properties of each neuronal circuit able to influence tauopathy spreading. In this study, we focused on a neuronal circuit well-known to efficiently propagate tauopathy and asked whether the stage of tau aggregation, in correlation with microglial activation, influences the propagation of tauopathy. We developed an *in vivo* model by injecting mice unilaterally in the dentate gyrus (DG) and studied the spreading of tau in the contralateral hippocampus where transduced neurons project.^39^ We used this model to analyze the influence of tau aggregation on its spreading by using three different AAVs allowing the expression of wild-type human tau (Tau^WT^), tau carrying the P301L mutation (Tau^P301L^), and an aggregation-prone variant co-expressing wild-type tau with a deltaK280 repeat-domain fragment (Tau^ProAgg^).^40^ We found greater spreading of human tau to the contralateral DG with aggregation-prone tau variants in 3-month-old C57BL/6J mice. We then applied the same paradigm to APP/PS1dE9 mice, in which amyloid plaques accumulate in the brain parenchyma; under the modified conditions required by this strain, no significant difference in spreading was detected between tau variants.

## MATERIAL & METHODS

### Animals

Mice were housed under 12-hour light/dark cycle with food and water *ad libitum*. Eight adult males C57Bl/6J mice (4-month-old, JANVIER laboratories) per group were used for AAV transduction. Eight adult females *APP/PS1dE9* mice (JAX MMRRC Stock# 034832) per group were used to assess the spreading of tau in an amyloidosis context and were compared to their WT littermates.

### Viral vectors design and production

In this study, we used two types of AAV backbones. First, to assess the relevance of using an AAV model to study tau spreading, we used a construct allowing the co-expression of GFP and the wild-type 1N4R isoform of human tau variants (Tau^WT^). A V5 tag was inserted between tau exons 3 and 4 in order to confidently distinguish the human tau protein of transgenic origin from the murine ortholog.^11,38,41^ A P2A sequence between the two transgenes allowed the cleavage of the polypeptide during the translation process, thus leading to an equimolar expression of the two proteins.^42^ Following AAV injection, this construct design ensures that only transduced cells expressed GFP, while the V5 tag was expressed in both transduced cells and recipient cells. The resulting transgene was subcloned into a shuttle plasmid for the production of AAV particles, under the control of the SYN2 promoter to restrict transgene expression to neurons. Second, to assess any correlation between tau aggregation status and spreading propensity, we compared three different tau variants based on the human 1N4R isoform. The first variant corresponded to the wild-type version (Tau^WT^), the second carried the P301L mutation (Tau^P301L^) which leads to familial forms of frontotemporal dementia, the third co-expressed the wild-type version with a peptide corresponding to tau repeat-domains bearing the dK280 mutation which promotes its aggregation (Tau^ProAgg^).^40^ These three variants were subcloned into a shuttle plasmid to produce AAV particles, under the control of the CBA promoter as described in d’Orange et al.^40^ A control vector based on the same backbone expressed GFP only. AAV production process is further detailed in *Supplementary materials & methods*.

### AAV stereotaxic injections

AAV injections were performed unilaterally with a stereotaxic frame to target the granular layer of the DG in the dorsal hippocampus, at the following coordinates: -2.5 mm posterior and +1.5 mm lateral relative to bregma, and -2.0 mm from the surface of the skull. After slowly lowering the injection canula at the right position, the AAV, diluted with PBS to obtain a volume of 2 µl, was injected over 10 minutes. The AAV dose depended on the experiment, with a dose of 5×10^10^ vector genome (vg) and 1×10^11^ vg for the first experiment (Fig. 1), of 5×10^9^ vg for the second set of experiments (Fig. 2-3-4) and 2.5×10^9^ vg for the last experiment (Fig. 5). A detailed procedure of the surgery and the AAV injection is provided in *Supplementary materials & methods*.

**Fig. 1.**
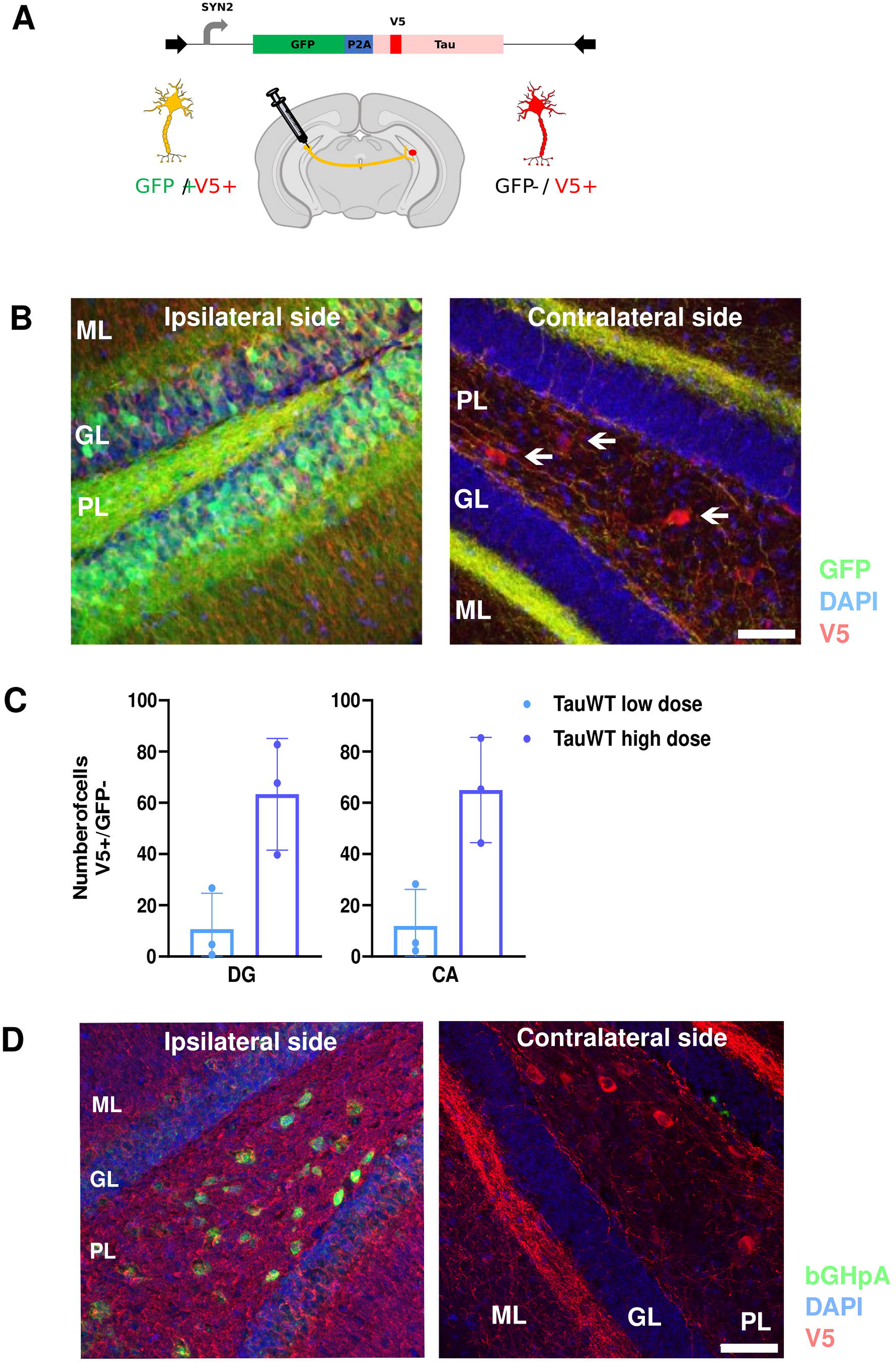
Proof of concept of the DG-targeted AAV model of tauopathy. **(A)** *Top*: schematic representation of the AAV vector design. Transgene expression is driven by the SYN2 promotor, followed by a GFP reporter gene, a P2A self-cleaving sequence, and the human tau gene carrying a V5 tag inserted between exon 3 and 4. *Bottom*: Schematic representation of the experimental approach. AAV is unilaterally injected into the DG to assess the spreading to contralateral DG. Neurons in the ipsilateral DG are expected to co-express GFP and V5, whereas contralateral DG neurons should express only V5. This diagram was created with BioRender. **(B)** Representative DG immunofluorescence images showing GFP (green), V5 (red), DAPI (blue) in the ipsilateral (left) and contralateral (right) DG. Arrows point to V5 positive neurons in the contralateral DG. PL: polymorphic layer; ML: molecular layer; GL: granule cell layer. Scale bar=50µm. **(C)** Quantification of V5-positive, GFP-negative neurons in the contralateral DG (left panel), and contralateral CA regions (right panel) after low-dose (blue) or high-dose (purple) AAV injection. Data are presented as mean ± SD; n=3 animals per group. **(D)** Immunofluorescence for V5 (red) and DAPI (blue) combined with *in situ* hybridization using a bGH polyA probe (green), specific for the viral genome. Scale bar=50µm.

**Fig. 2.**
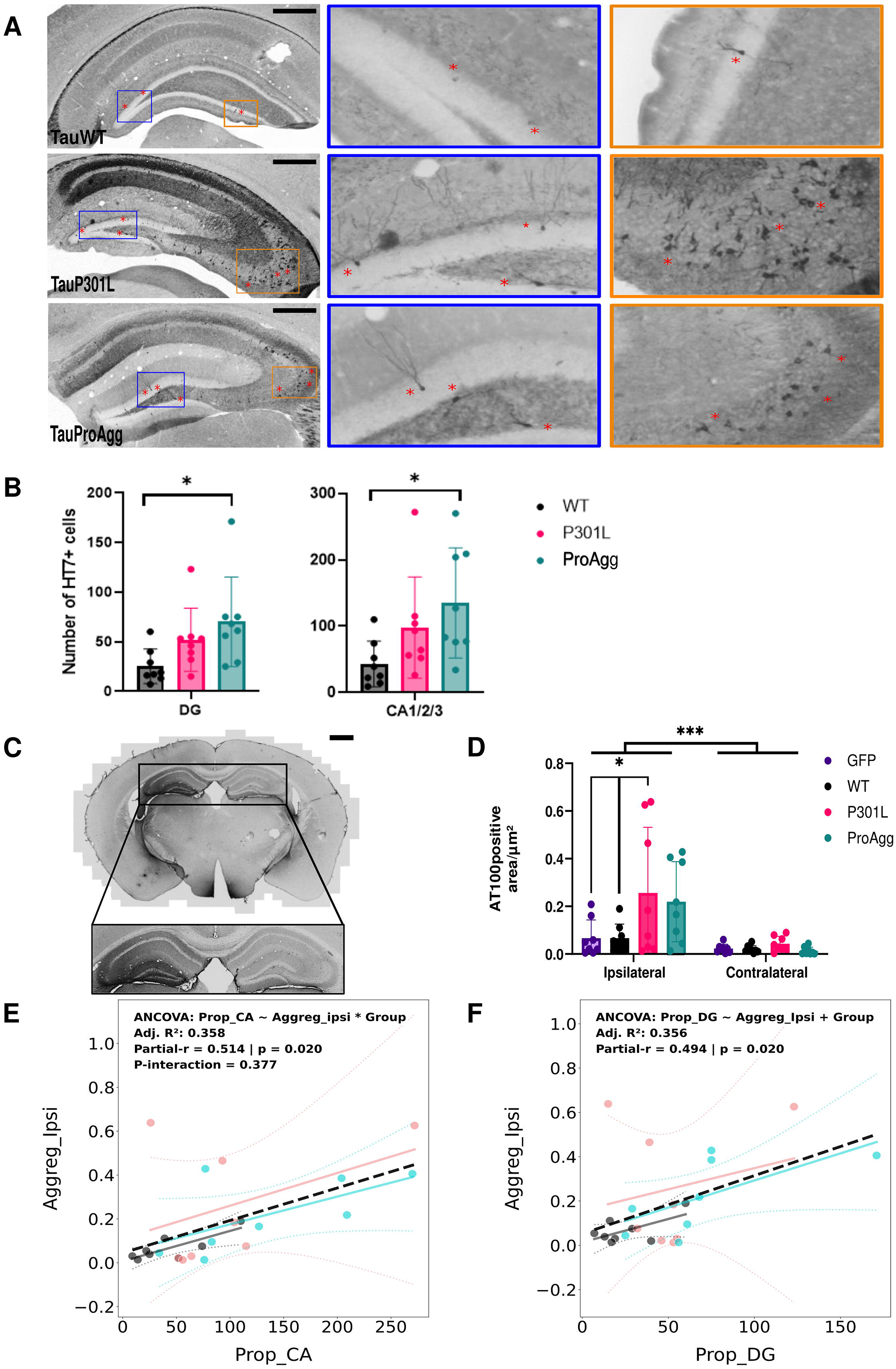
Impact of tau aggregation state on trans-synaptic spreading in the contralateral hippocampus and characterization of tau aggregation and neurodegeneration. **(A)** Representative HT7 immunohistochemistry images acquired at 20x using epifluorescence microscopy in Tau^WT^ (top), Tau^P301L^ (middle), and Tau^ProAgg^ (bottom) mice. Zooms of the DG polymorphic layer (blue boxes) and CA3 region (orange boxes) show that tau expression is dependent on the protein aggregation status. Red asterisks highlight some HT7 positive neurons. Scale bars=200 µm. **(B)** Quantification of HT7 positive neurons in the contralateral DG (left panel) and contralateral CA1/2/3 regions (right panel) for Tau^WT^ (black), Tau^P301L^ (pink), and Tau^ProAgg^ (cyan) groups. Data are presented as mean ± SD, n=8/group. Kruskall-Wallis test followed by Dunn’s multiple comparisons test; * p<0.05. **(C)** Representative AT100 immunohistochemistry image (10x) of a Tau^ProAgg^ injected brain slice (ipsilateral side shown on the left). Scale bar=200 µm. Zoom of the hippocampus in the black box below. **(D)** Quantification of tau aggregates density in ipsilateral and contralateral hippocampus. **(E-F)** Scatter plots showing the correlation of tau aggregation in ipsilateral hippocampus with respectively tau spreading in contralateral CA (E; Statistical significance was assessed by ANCOVA [Group x Aggregation interaction, *p<0.05]) and DG (F; Data were analyzed using ANCOVA with a common slope model [non-significant interaction removed; p=0.163]; Partial R values indicate the correlation between ipsilateral tau aggregation and DG propagation across all groups [*p < 0.05]), for WT mice injected with Tau^WT^ (black), Tau^P301L^ (pink) or Tau^ProAgg^ (cyan) AAVs. Solid lines represent linear regressions for each group; black dotted line represents global correlation.

**Fig. 3.**
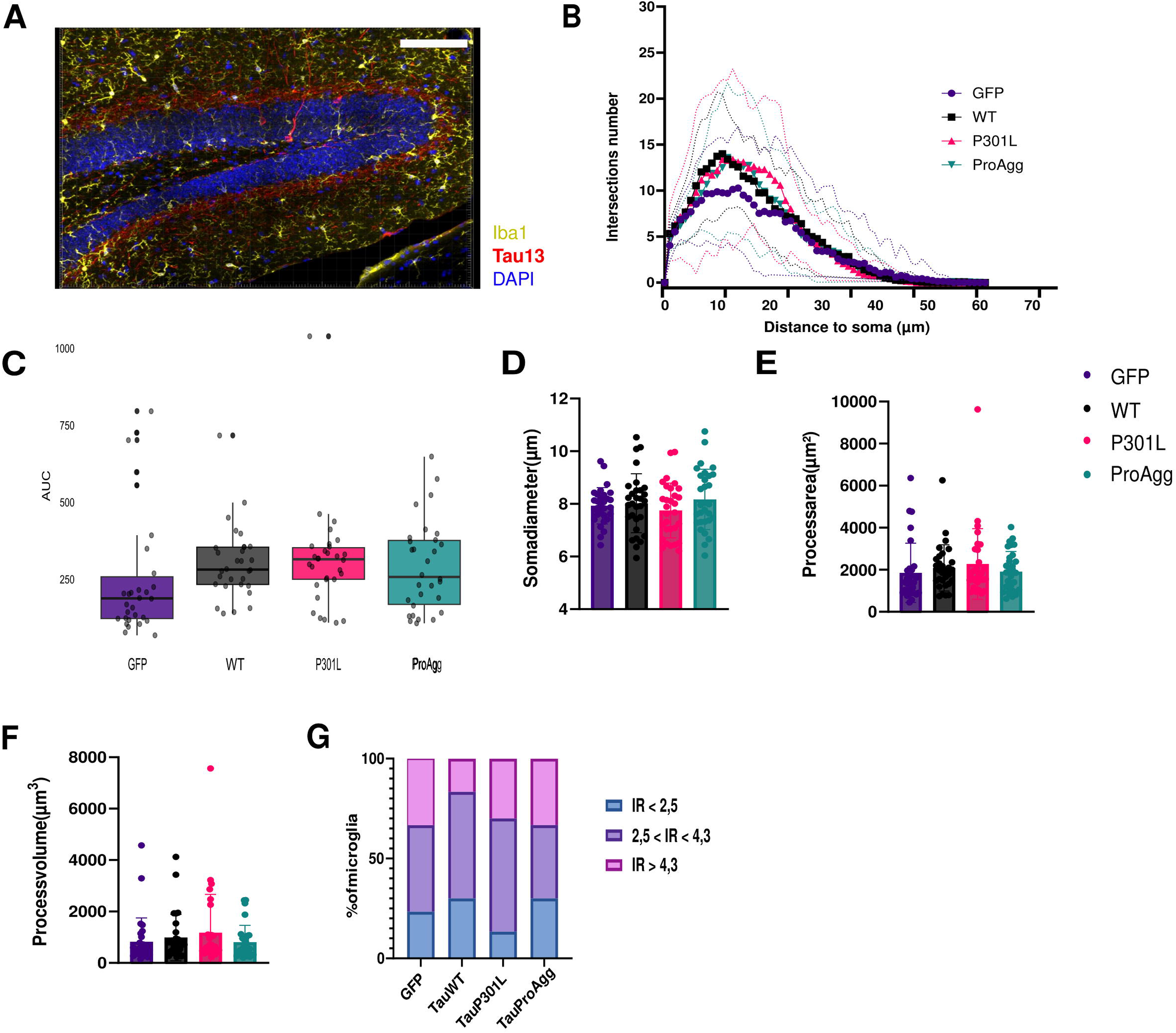
Characterization of microglial inflammation in the DG model. **(A)** Representative immunofluorescence of Tau13 (red), Iba1 (yellow) and DAPI (blue) showing the region analyzed for microglial morphology. Scale bar =200µm. **(B)** Sholl analysis of microglial morphology at vicinity of tau spreading site in the contralateral DG, showing the number of intersections as a function distance from the soma. Statistical analysis was performed using a linear mixed-effects model (LMM) to account for the hierarchical structure of the data (nested random effects: cells nested within images, nested within animals). The model included Group, Radius, and their interaction as fixed effects. P-values for the fixed effects were estimated using Satterthwaite’s approximation. Post-hoc pairwise comparisons were conducted point-by-point at each radius using Tukey’s HSD adjustment. No significant differences were observed between groups at any specific radii (p > 0.05)]. Mean ± SEM; n=5 cells from 2 fields of view from 3 animals per group. **(C-F)** Microglial morphological parameters: (C) Area under the curve (AUC) from Sholl curves, (D) soma diameter (µm), (E) ramification area (µm²), (F) ramification volume (µm³). Mean ± SD; n=30 microglia/group and 3 animals/group; linear mixed-effects model using Satterthwaite’s method, p-values adjusted using Tukey’s post-hoc test for multiple comparisons. **(G)** Repartition of microglia in function of the IR (IR=Sholl peak / number of primary intersections). Chi2 test.

### Histology and immunolabeling

To collect brains for histology, mice were injected with buprenorphine (0.3 mg/kg) 30 minutes before euthanasia with a lethal dose of ketamine (220 mg/kg) and medetomidine (0.7 mg/kg). Blood vessels were then washed from blood cells by trans-cardiac perfusion with 10 mL of 0.9% NaCl solution, followed by tissue fixation with 100 mL of 4% paraformaldehyde (PFA) in phosphate-buffered saline (PBS). Whole brains were extracted from the skull and post-fixed in 4% PFA for 24 h before being immersed in PBS containing 30% sucrose for cryoprotection and stored at 4°C. Brains were then cut as 30 or 40 µm-thick free-floating coronal sections in 10 series with a freezing microtome and stored in cryoprotectant solution at -20°C until further processing.

All the immunostaining process was performed on free-floating sections, which were successively rinsed in PBS, incubated in blocking solution (PBS1X containing 5% normal goat serum and 0.2% Triton X-100) for 1 h at room temperature before being transferred to primary antibody solution for at least one overnight incubation at 4°C. For immunofluorescence staining, sections were incubated with Alexa Fluor-conjugated secondary antibody (ThermoFisher). Sections were finally incubated in DAPI solution for 5min and mounted on slides under coverslips with Mowiol. For immunohistochemistry, sections were incubated with biotinylated secondary antibody, then incubated 1h with Avidine-Biotine complex (Kit ABC, VectorLab SK-6100) and with DAB solution (VectorLab SK-4100) until conversion of the chromogenic substrate. Sections were rinsed with 0.01M PBS between each step. Brain sections were then mounted on slides and dehydrated with increasing concentration ethanol solutions and xylene before mounting in Eukitt (Sigma 03989). All references, as well as final concentration of antibodies are indicated in Supp. Table 1 of the *Supplementary materials & methods*.

We also performed *in situ* hybridization (ISH) using a riboprobe designed specifically to detect the AAV recombinant genome in the transduced cells. This riboprobe was specific for the bovine Growth Hormone polyadenylation signal inserted in the AAV expression cassette. ISH was coupled with V5 (Fig. 1D) or Tau13 (Suppl. Fig. 1A and Suppl. Fig. 2E) immunostainings. The protocol is detailed in *Supplementary materials & methods*.

### Image Analysis

Quantification of tauopathy spreading following AAV-Tau injection was performed by manual counting of labelled cells under an epifluorescence microscope (Leica DM6000B). This manual quantification was done by screening all the brain slices covering the whole antero-posterior axis of the hippocampus (i.e. from -0.9 to -4 mm relative to bregma) in 1 series for each animal after immunolabelling with V5, HT7 or Tau13 antibodies. During quantification contralateral DG and contralateral cornu ammonis (CA) were analyzed as distinct regions. Manual counting was performed blind to the experimenter with respect to experimental group. Samples were labeled with the animal’s identification number assigned to all individuals of the strain within the animal facility, based on birth order or, where applicable, at the time of genotyping. The experimental group was recorded in the injection report to ensure traceability.

Quantitative analysis of AT100 (tau aggregation), NeuN (neurodegeneration) and BAM10 (density of amyloid plaques) immunostainings were made on digitized images of the brain sections using a slide-scanner (Zeiss Axio Scan Z1) at x20 magnification. Analysis was conducted on all brain slices of one series from – 2.5 mm to +3.5 mm relative to bregma for each animal. The method used to quantify those parameters is exposed in detail in *Supplementary materials & methods*.

The morphology of microglia was assessed on confocal views with IMARIS 9 software. Before the analysis, we acquired images using a spinning disk confocal microscope (Nikon CSU-W1), at x63 magnification, with mosaic and a stack set to acquire 30µm of thickness with a z-step of 0.3µm. We chose to reconstruct and analyze microglia in the contralateral molecular layer of the DG, close to the granular layer, due to its high density of transgenic tau-positive fibers from mossy cells transduced in the ipsilateral DG and projecting in this area. We analyzed 5 microglia per brain section, with 2 sections per animal and 3 animals per group, i.e. 30 cells in total per group. After clipping the individual microglial cells based on Iba1 labeling, IMARIS 9 allowed to extract different parameters such as Sholl analysis, soma diameter, ramification index, sum filament area and filament volume. Sholl circles were set at 1µm apart. From Sholl analysis, we extracted the area under the curve parameter.

### Isolation and characterization of brain-derived extracellular vesicles

In order to isolate mouse brain-derived extracellular vesicles (EVs), we injected 8 WT mice bilaterally with either Tau^WT^ or Tau^ProAgg^, while 2 uninjected mice served as controls. After 4 weeks, mice were euthanized, their brain was extracted, and their hippocampi and cortex were dissected out and kept frozen at -70°C. EVs were isolated as described below from the pooled hippocampi and cortices of 2 mice, generating 4 EV preparations for each condition (Tau^WT^, samples H2 to H5; Tau^ProAgg^, samples H6 to H9) and 1 control EV preparation (sample H1). The protocol is detailed in *Supplementary materials & methods*. Briefly, tissues were gently dissociated in a specific buffer with Collagenase D in a thermomixer. Tissue debris were removed by centrifugation and filtration and EVs purified by size-exclusion chromatography and stored at -80°C until use. EVs were characterized by negative-staining electron microscopy (EM) and immunogold analysis, as described before (see *Supplementary materials & methods* for details). ^43,44^ For immunoblotting EVs were spotted on nitrocellulose membranes as described in Supplementary materials. CD9/63/81 antibodies were used to characterize the EVs, and AT8 antibody to assess their tau cargo. All references, as well as final concentration of antibodies are indicated in Supp. Table 1.

### Statistical analysis

Methodology for statistical analyses is directly reported in the results section as well as in the figure’s legends. These analyses were carried out using GraphPad Prism version 9.1 (GraphPad® software, San Diego, CA, USA) with two exceptions performed with RStudio:

- Sholl analysis (Fig. 3B and 5A) and microglia morphological parameters (Fig. 3C-F and Fig. 5 B-E) were analyzed with a linear mixed-effects model (LMM) with Group, Radius and their interaction included as fixed effects and the number of intersections was defined as the dependent variable. A nested random-intercept structure (microglia nested within scan nested within each individual animal) to account for hierarchical data dependencies. This model was implemented using lme4 and lmerTest packages in R. P-values for fixed effects were estimated using Satterthwaite’s approximation for degrees of freedom. Post-hoc pairwise comparisons between groups were performed using the emmeans package with Tukey’s adjustment for multiple comparisons. To further investigate the spatial distribution of microglial complexity, post-hoc pairwise comparisons were performed at specific distances from the soma (every 4 µm) using the emmeans package. These local contrasts were derived from the linear mixed-effects model to identify the specific radii where the groups significantly diverged. P-values were adjusted for multiple comparisons using Tukey’s HSD (Honest Significant Difference) method to maintain a family-wise error rate of α= 0.05.
- To evaluate the relationships between various parameters and Tau propagation (e.g., Soma Diameter vs. CA propagation), Analysis of Covariance (ANCOVA) was performed. The initial model included Group, the covariate (Y), and their interaction as independent variables to assess potential differences in regression slopes. Following standard procedures for model parsimony, non-significant interaction terms (p > 0.05) were excluded from the final analysis to evaluate the main effects of Group and Y. Partial correlation coefficients (Partial_R) were calculated to determine the specific association between Y and X, accounting for group-related variance (Fig. 2E-F; Fig. 4G-L; Fig. 5G).

**Fig. 4.**
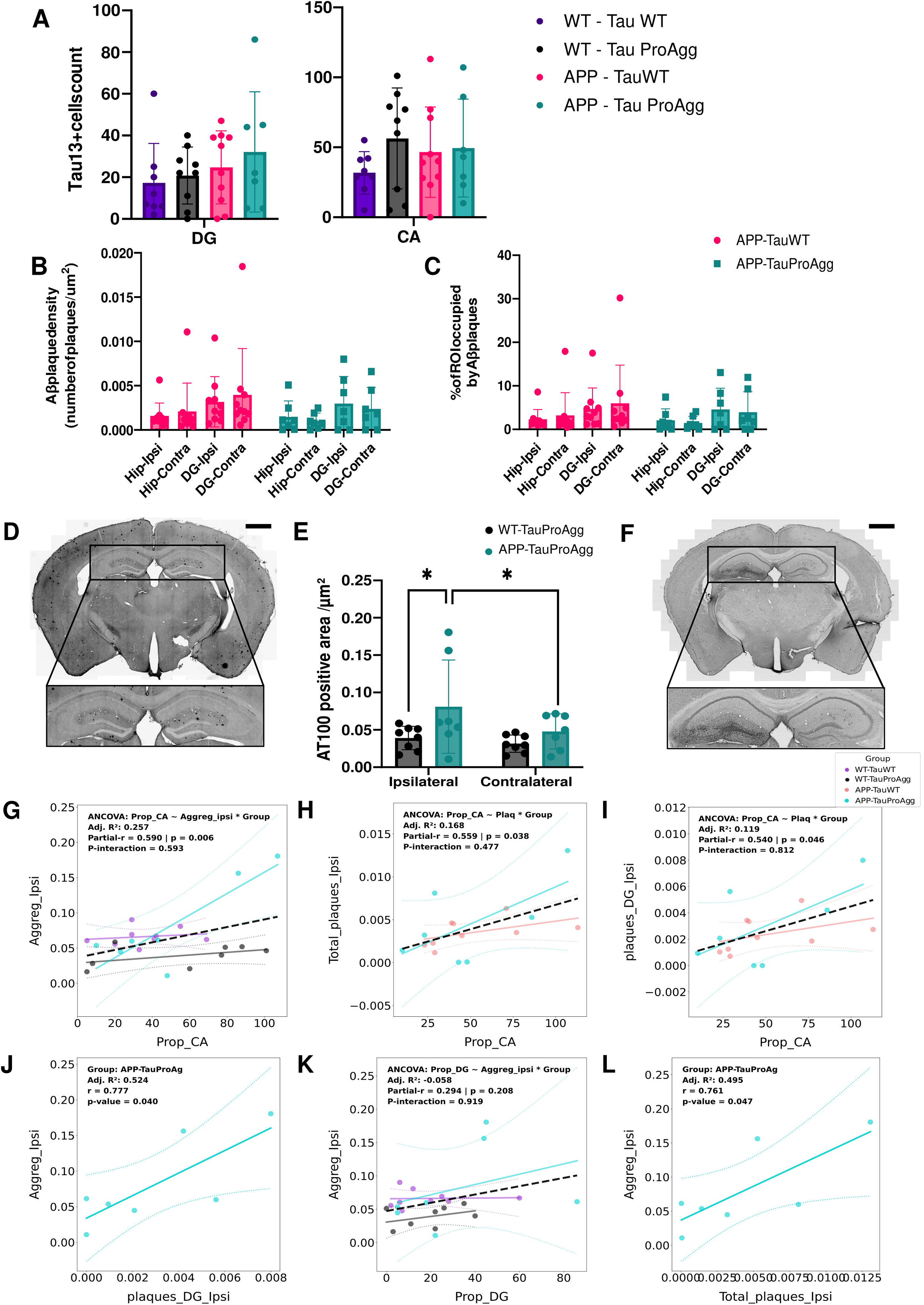
Influence of tau aggregation and amyloidosis on tau spreading in an amyloidosis context. (A) Quantification of Tau13 positive neurons in contralateral DG (left) and CA1/2/3 region (right), in WT mice injected with Tau^WT^ (purple) or Tau^ProAgg^ (black), and APP/PS1 mice injected with Tau^WT^ (pink) or Tau^ProAgg^ (cyan). Mean ± SD; n=8/group; Kruskall-Wallis test followed by Dunn’s multiple comparisons test. (B-C) Quantification of Aβ plaques in the hippocampus and DG (ipsilateral and contralateral) in APP/PS1dE9 mice injected with Tau^WT^ (pink) or Tau^ProAgg^ (cyan), measuring plaque number (B) and size (C) relative to ROI surface. Mean ± SD; n=8/group; Two-ways ANOVA with Sídák’s multiple-comparisons test; * p <0.05, ** p< 0.01. (D) Representative BAM10 immunohistochemistry of an AAV-Tau^ProAgg^ injected mouse brain slice (ipsilateral side shown on the left). Scale bar=200µm. Zoom of the hippocampus in the black box below. (E) Quantification of tau aggregates density in ipsilateral and contralateral hippocampus. Mean ± SD; n=8/group. Two-way ANOVA followed by the two planned comparisons described in the Methods (Fisher’s LSD, unadjusted); *p<0.05; exact p-values are given in the Results. (F) Representative AT100 immunohistochemistry of an AAV-Tau^ProAgg^ injected mouse brain slice (ipsilateral side shown on the left). Scale bar=200µm. Zoom of the hippocampus in the black box below. (G-L) Correlations between tau spreading in CA and tau aggregation in ipsilateral hippocampus (G; Statistical significance was assessed by ANCOVA [Group x Aggregation interaction], **p<0.01), amyloid plaques density in the whole ipsilateral hippocampus (H; Statistical significance was assessed by ANCOVA [Group x Amyloidosis interaction], *p<0.05), and amyloid plaques density in ipsilateral DG (I; Statistical significance was assessed by ANCOVA [Group x Amyloidosis interaction], *p<0.05). (J) Correlation between amyloid plaque density in ipsilateral DG and tau aggregation in ipsilateral hippocampus (Pearson’s test). (K) Correlation between tau spreading in contralateral DG and tau aggregation in ipsilateral hippocampus (ANCOVA [Group x Aggregation interaction], p=0.208). (L) Correlation between amyloid plaque density in the whole ipsilateral hippocampus and tau aggregation in ipsilateral hippocampus (Pearson’s test); respectively in purple/black/pink/cyan, WT-Tau^WT^ / WT-Tau^ProAgg^ / APP-Tau^WT^ / APP-Tau^ProAgg^.

## RESULTS

### AAV-mediated overexpression of human tau in the mouse dentate gyrus to study tauopathy spreading

To investigate the spreading of human tau protein between connected neurons, we developed a model based on AAV-mediated gene transfer targeted to the dentate gyrus (DG) of adult mice. The AAV vector used expresses two transgenes, GFP and the 1N4R human tau isoform under the control of the neuron-specific SYN2 promoter. A self-cleaving P2A sequence was included between the two transgenes to allow independent translation of the two proteins (Fig. 1A). To enable detection of transgenic tau, a V5-tag was inserted between exons 3 and 4 of the tau cDNA. Stereotaxic injections were performed unilaterally in the dorsal DG, targeting both granule cells and mossy cells, the latter of which being known to project to the contralateral DG.^39^ This anatomical feature enables a spatial distinction between neurons directly transduced by the AAV vector and neurons potentially receiving transgenic tau via trans-synaptic spreading.

In the ipsilateral DG, neurons exhibited co-expression of GFP and V5-tagged tau (Fig. 1B left panel). In contrast, in the contralateral DG, some neurons were V5-positive but GFP-negative (Fig. 1B right panel), indicating the presence of transgenic tau in the absence of AAV transduction. Notably, labeled axon fibers, either GFP-positive alone or co-labeled with tau, were also observed in the contralateral DG, corresponding to mossy cells projections from the injection site.

A dose-response pilot study was conducted to determine the optimal AAV dose that maximized neuronal transduction while minimizing toxicity. Quantification of V5-positive neurons in the contralateral hippocampus (by manual counting distinguishing DG and CA subregions) showed a clear dose-dependent increase in tau spreading with higher AAV doses (Fig. 1C). To exclude the possibility that tau expression in the contralateral hippocampus was due to vector diffusion, we performed *in situ* hybridization using a probe targeting the bGHpA sequence of the AAV recombinant genome, combined with V5 immunostaining (Fig. 1D). In the ipsilateral DG, cells co-expressed bGHpA signal and V5 (Fig. 1D, left panel), whereas in the contralateral DG, V5-positive neurons lacked viral genome signal (Fig. 1D, right panel). These findings confirm that tau detection in the contralateral DG results from protein spreading, and not from inadvertent viral transduction.

Altogether, this experimental paradigm provides a robust and anatomically-defined system to study tau propagation *in vivo*.

### Influence of tau aggregation on its propagation

Preliminary experiments (data not shown) showed that vectors containing the GFP reporter gene and the V5 tag did not result in a significant level of tau aggregation unlike our previously described constructs^40^ free of GFP, P2A sequence and V5 tag. We therefore used these to investigate whether tau aggregation influenced its propagation. We took advantage of three different AAVs, each expressing one of three variants of the human 1N4R tau isoform: wild-type (AAV-Tau^WT^), P301L mutant (AAV-Tau^P301L^) and a highly aggregation prone version co-expressing the wild-type tau and the deltaK280-mutated tau repeat-domain (AAV-Tau^ProAgg^).^40^ To assess how these tau species impact propagation, we injected 3-month-old C57BL/6J male mice and counted HT7-positive neurons in the contralateral DG. To compensate for the high activity of the CBA promoter present in these constructs, compared to the SYN promoter used for the experiments depicted in figure 1, we decreased the vector dose down to 5×10^9^ vg. We observed tau spreading to the contralateral hippocampus with all three different forms of tau (Fig. 2A). Notably, a significant increase of HT7-positive neurons was detected in the contralateral DG (Fig. 2B, Kruskal-Wallis test, H(2)=8.51; *p<0.05; N=24; Post-hoc comparisons using Dunn’s test; Z=2.901; adj.p=0.0112) and CA (Fig. 2B, Kruskal-Wallis test, H(2)=8.12; *p<0.05; N=24; Post-hoc comparisons using Dunn’s test; Z=2.776; adj.p=0.0165) regions of mice injected with Tau^ProAgg^ AAVs, with a similar trend observed for Tau^P301L^ (Fig. 2B).

To ensure that the contralateral HT7 signal reflected tau propagation rather than vector diffusion, we performed *in situ* hybridization combined with Tau13 immunostaining. While the AAV specific probe was restricted to the ipsilateral DG, Tau13 immunoreactivity was present in both ipsilateral and contralateral hippocampi (Suppl. Fig. 1A).

To evaluate the relationship between aggregation level and propagation, we performed an immunolabelling with AT100 (Fig. 2C), which is specific for abnormal conformations of tau, and compared signal intensities between ipsilateral and contralateral hippocampi across all groups. We observed an increase of AT100 labelled area in the ipsilateral hippocampus of Tau^P301L^-injected mice compared to Tau^WT^ and GFP groups (Fig. 2D, two factors ANOVA, Post-hoc analysis using Sidak’s multiple comparisons test, Tau^P301L^ values versus GFP [t(56)=3.18, *p<0.05] and Tau^WT^ [t(56)=3.17, *p<0.05] groups). Although not statistically significant, Tau^ProAgg^ also showed a trend toward increased aggregation in the injected hippocampus relative to Tau^WT^ and GFP groups (Fig. 2D, Post-hoc analysis using Sidak’s multiple comparisons test, Tau^ProAgg^ values versus GFP group [t(56)=2.557, p=0.0772] and Tau^WT^ [t(56)=2.549, p=0.0788]). Across all conditions, AT100 labeling was higher in the injected hippocampus compared to the contralateral one (Fig. 2D, two-way ANOVA, Group [F(3,28)=3.13, *p<0.05] and the Ipsi/Contra factor [F(1,28)=18.32, ***p<0.001]). Notably, AT100 immunoreactivity did not differ significantly between Tau^P301L^ and Tau^ProAgg^, although Tau^ProAgg^ produced greater contralateral signal. AT100 labelling therefore does not by itself account for the difference in spreading between these two variants. One possibility is that the two constructs generate conformationally distinct assemblies that AT100 detects with comparable efficiency but that differ in their capacity to be released and taken up; this remains to be tested.

To determine whether tau spreading is related to the severity of cell death, we quantified tissue volume in four hippocampal subregions - the whole hippocampus, the whole DG, the polymorphic layer of the DG and the CA1 layer - extrapolated from serial section areas (Suppl. Fig. 1 I-L).

In all three tau groups, the ipsilateral side was smaller than the contralateral side for the whole hippocampus (Suppl. Fig. 1I; two-way ANOVA, Ipsi/Contra [F(1,22)=13.40, **p<0.01], Ipsi/Contra x Group [F(3,22)=5.812, **p<0.01]; Sidak post-hoc, Tau^WT^ [t(22)=3.242, **p<0.01], Tau^P301L^ [t(22)=2.406, *p<0.05], Tau^ProAgg^ [t(22)=4.567, ***p<0.001]) and for the whole DG (Suppl. Fig. 1J; Ipsi/Contra [F(1,22)=22.98, ****p<0.0001], Ipsi/Contra x Group [F(3,22)=8.154, ***p<0.001]; Sidak post-hoc, Tau^WT^ [t(22)=3.162, **p<0.01], Tau^P301L^ [t(22)=2.862, **p<0.01], Tau^ProAgg^ [t(22)=6.311, ****p<0.0001]). In the polymorphic layer, the same ipsilateral reduction was restricted to the Tau^P301L^ and Tau^ProAgg^ groups (Suppl. Fig. 1L; Group [F(3,22)=3.802, *p<0.05], Ipsi/Contra [F(1,22)=17.84, ***p<0.001]; Sidak post-hoc, Tau^P301L^ [t(22)=3.229, **p<0.01], Tau^ProAgg^ [t(22)=2.794, *p<0.05]). Across regions, the largest ipsilateral reduction was consistently observed in the Tau^ProAgg^ group. In addition, ipsilateral DG volume was smaller in Tau^ProAgg^ mice than in GFP and Tau^P301L^ mice (Suppl. Fig. 1J; Sidak post-hoc, versus GFP [t(44)=4.058, **p<0.01], versus Tau^P301L^ [t(44)=3.543, **p<0.01]), consistent with the atrophy described above. In CA1, the opposite was found: both ipsilateral and contralateral volumes were larger in Tau^ProAgg^ mice than in GFP and Tau^P301L^ mice (Suppl. Fig. 1K; Group [F(3,22)=5.064, **p<0.01], Subject [F(22,22)=6.755, ****p<0.0001]; Sidak post-hoc, ipsilateral versus GFP [t(44)=2.990, *p<0.05] and versus Tau^P301L^ [t(44)=3.319, *p<0.05]; contralateral versus GFP [t(44)=2.924, *p<0.05] and versus Tau^P301L^ [t(44)=2.917, *p<0.05]). Tau^WT^ mice differed from Tau^ProAgg^ mice in CA1 on the contralateral side only, where their volume was smaller (Suppl. Fig. 1K; Sidak post-hoc, [t(44)=2.794, *p<0.05]). Because the CA1 difference was present on both sides, it is unlikely to result from the unilateral injection and may instead reflect inter-animal variability in the extrapolation of volumes from serial sections. In summary, Two distinct effects were apparent: a reduction in volume on the ipsilateral side compared to the contralateral side within the group, and differences in overall volume between the groups.

Given the suspected role of microglia in mediating synaptic tau propagation, we characterized their morphology in the contralateral hippocampus, near the axonal projections of transduced mossy cells, an area highly enriched in tau-positive fibers projecting from the ipsilateral hippocampus (Fig. 3A). We analyzed the 3D morphology of 10 microglia per animal (5 microglia on 2 different fields of view) in 3 animals per group.

Microglial morphology did not differ significantly between groups. Sholl analysis revealed a significant main effect of Radius [F(1,7316)=5009.46, p<0.001] and a significant Group x Radius interaction [F(3,7316)=17.99, p<0.001], indicating non-parallel branching profiles, but no main effect of Group [F(3,8.5)=1.19, p=0.369] (Fig. 3B; linear mixed-effects model with Satterthwaite’s method). Post-hoc pairwise comparisons (Tukey-adjusted) at individual radii (0 to 60 um) did not identify any point of divergence between GFP, Tau^WT^, Tau^P301L^ and Tau^ProAgg^ groups (all p>0.05). The significant interaction therefore reflects a difference in the shape of the branching profiles that is not localised to any specific distance from the soma once corrected for multiple comparisons.

No other morphological parameter differed between groups: Sholl AUC (Fig. 3C; [F(3,7.99)=0.15, p=0.93]), soma diameter (Fig. 3D; [F(3,40.7)=0.75, p=0.53]), process area (Fig. 3E; [F(3,73.49)=0.15, p=0.93]) and process volume (Fig. 3F; [F(3,111.02)=0.34, p=0.8]); all assessed with linear mixed-effects models with Satterthwaite’s method. The distribution of microglia across ramification classes was likewise unchanged (Fig. 3G; chi-square, X2=5.907, df=6, p=0.4337), although a higher proportion of highly ramified cells was apparent in the GFP and Tau^P301L^ groups.

Finally, correlation analysis revealed that tau aggregation in the ipsilateral hippocampus (quantified based on AT100 Immunolabelling) positively correlated with propagation to the contralateral CA (Fig. 2E, ANCOVA, Partial-r=0.514; *p< 0.05) and DG (Fig. 2F, ANCOVA, Partial-r=0.494, *p< 0.05; the interaction term was not significant and thus excluded [p-value=0.163]) regions of the contralateral hippocampus in tau-injected animals.

### The aggregation-prone form of tau is more efficiently loaded within extracellular vesicles (EVs)

Extracellular vesicles (EVs) might play an important role in neuron-to-neuron spreading of pathological forms of tau. Moreover, recent literature highlighted a close link between microglia and tau spreading but the mechanisms are still unknown.^12,24^ In fact, the current hypothesis suggests that microglia are able to internalize tau aggregates produced by neurons, partially degrade them and release tau seeds within EVs, thereby contributing to tau spreading.^12,35,36^ To assess whether EVs carry tau proteins in our model, we isolated EVs from hippocampal and cortical tissues of mice after injection in the DG of Tau^WT^ or Tau^ProAgg^ AAVs. Electron microscopy analysis of EV preparations revealed no significant differences in their shape or morphology, and the size distribution of EVs was comparable between samples with a median vesicle diameter of ∼80±7 nm (Suppl. Fig. 1B-C and data not shown). Immunoblotting experiments revealed that, as expected, EV preparations contained comparable amounts of the CD9, CD63 and CD81 EV markers (Suppl. Fig. 1D-E). Interestingly, AT8-positive tau was detected to various extents in hippocampal EVs from AAV-Tau^ProAgg^-injected mice but not in AAV-Tau^WT^ ones (Suppl. Fig. 1F).

To confirm these results, we performed a semi-quantitative immunogold-labelling EM assay on the EV preparations and counted CD9/63/81-positive EVs on one hand and AT8-positive EVs on the other (Suppl. Fig. 1H). The number of CD9/63/81-positive EVs was comparable in Tau^WT^- and Tau^ProAgg^-injected mice, whereas AT8-positive hippocampal EVs were markedly more numerous in Tau^ProAgg^-injected mice (Suppl. Fig. 1G; two-way ANOVA, labelling factor [F(1,6)=56.01, ***p<0.001]; Sidak post-hoc, AT8 labelling in Tau^WT^ versus Tau^ProAgg^ [t(12)=3.129, *p<0.05]). These immunogold data agree with the immunoblotting results. AT8-positive EVs were not detected in preparations from hippocampus of non-injected animals, nor in cortex from animals injected in the hippocampus (data not shown).

Together, these experiments establish that AT8-positive tau is associated with EV preparations from Tau^ProAgg^-injected mice and is more abundant there than in preparations from Tau^WT^ injected-mice. EVs therefore represent a plausible vehicle for the dissemination of pathological tau in this model. Determining whether this EV-associated tau is intraluminal and seed-competent, and whether EVs are required for the spreading measured in the previous experiments are the natural next steps.

### Influence of amyloidosis on tau spreading

As literature showing an influence of amyloidosis on tauopathy progression continues to grow, it became essential to assess its influence on the synaptic spreading of pathological tau. Two groups of mice – APP/PS1dE9 and WT– were injected with AAVs encoding either Tau^WT^ or Tau^ProAgg^. We first observed a surprisingly high mortality rate following AAV-Tau injection in APP/PS1dE9. These transgenic mice are indeed very fragile around the age of 3 months, and we had to adapt our experimental protocol. Consequently, we chose to inject 5-month–old female mice with a lower dose than in the previous experiment and to study the spreading 1 month post injection. In these conditions, we did not observe any statistically significant difference in tau spreading between the four groups but there were slightly more Tau13-positive cells in the contralateral DG of APP mice than in WT mice, and slightly more in Tau^ProAgg^-injected mice than in Tau^WT^-injected ones in both WT and APP mice (Fig. 4A). Additionally, we identified Tau13-positive neurons in several other contralateral regions including the retrosplenial cortex, entorhinal cortex, other cortical regions and subiculum (Suppl. Fig. 2A-D). To confirm that Tau13-positive cells in the contralateral hippocampus resulted from tau spreading rather than viral diffusion, we performed an *in situ* hybridization as described above. As expected, bGH signal was restricted to the injection site, but not visible in the contralateral side, while Tau13 labelling was evident on both ipsilateral and contralateral sides (Suppl. Fig. 2E).

We next assessed amyloid pathology in APP/PS1dE9 mice injected with either Tau^WT^ or Tau^ProAgg^ AAVs (Fig. 4B-C; Representative BAM10 immunostaining is shown in Fig. 4D). We quantified both the number of Aβ plaques and the plaque-covered area in the DG and CA1–3 regions, relative to total hippocampal area, on both sides. No significant differences were found between APP/PS1dE9 mice injected with Tau^WT^ versus Tau^ProAgg^ AAVs.

We then analyzed tau aggregation by AT100 immunohistochemistry, using the same protocol as before (Fig. 4E-F). Both pre-specified hypotheses were supported: tau aggregation was higher in the ipsilateral hippocampus of AAV-Tau^ProAgg^-injected APP mice than in their WT counterparts (Fig. 4E; two-way ANOVA with uncorrected Fisher’s LSD, [t(26)=2.41, *p=0.023]), and higher on the ipsilateral than on the contralateral side in APP mice (Fig. 4E; [t(13)=2.18, *p=0.048]). Both p-values are unadjusted, as these were comparisons specified a priori. As previously reported, no tau aggregates were detected in Tau^WT^-injected mice. Aggregation in these AAV-Tau^ProAgg^-injected mice was on average about two times lower than in the animals of the first experiment (Fig. 2D), which we attribute to the lower AAV dose and the shorter post-injection interval.

We then examined how spreading, tau aggregation and amyloid load relate to one another. Tau spreading to the CA region correlated positively with ipsilateral tau aggregation (Fig. 4G; ANCOVA, Partial-r=0.59, **p<0.01), whereas spreading to the DG did not (Fig. 4K; ANCOVA, Partial-r=0.294, p=0.208). Spreading to the CA region also correlated with amyloid plaque load, both in the whole ipsilateral hippocampus and in the ipsilateral DG (Fig. 4H, I; ANCOVA, Partial-r=0.559, *p<0.05 and Partial-r=0.54, *p<0.05). Finally, in Tau^ProAgg^-injected APP mice, amyloid plaque load correlated with tau aggregation in the ipsilateral DG and in the whole ipsilateral hippocampus (Fig. 4J, L; r=0.777, *p<0.05 and r=0.761, *p<0.05).

We also assessed the extent of neurodegeneration in our model. As in our prior experiment, we focused on the whole hippocampus (Suppl. Fig. 2F), the DG (Suppl. Fig. 2 G), the CA1 region (Suppl. Fig. 2H), and the polymorphic layer of the DG (Suppl. Fig. 2I) on both ipsilateral and contralateral sides. As a proxy of neurodegeneration, the volume of each structure was extrapolated from surface areas measured on a series of brain sections. We did not observe any difference between groups and between ipsilateral and contralateral sides for any of the regions examined.

Next, we analyzed microglial morphology using the above-described method. A significant morphological shift was detected in the hippocampus of WT mice injected with AAV-Tau^ProAgg^ compared to the other groups (Fig. 5A; linear mixed model using Satterthwaite’s method, p-values adjusted using Tukey’s correction). This was illustrated by the Sholl statistical analysis using a linear mixed-effects model analysis which revealed a significant effect of Group [F(3, 10.5) = 5.74, p = 0.014] and Radius [F(1, 7556) = 6348.16, p < 0.001]. A significant Group x Radius interaction was observed [F(3, 7556) = 19.89, p < 0.001], indicating that differences in microglial branching between groups were dependent on the distance from the cell body. Post-hoc pairwise comparisons (Tukey-adjusted) further confirmed these differences. At proximal distances from the soma (0–16 µm), microglia exhibited significantly higher branch complexity in both Tau^WT^ and Tau^ProAgg^-injected APP mice compared to Tau^ProAgg^ -injected WT mice’s ones (e.g., at 0 µm: p = 0.002 and p = 0.004, respectively). Furthermore, microglia from Tau^ProAgg^ -injected WT mice showed a significant reduction in intersections compared to Tau^WT^-injected WT mice at these same proximal distances (0 µm: p = 0.019; 4 µm: p = 0.033). These morphological differences gradually diminished at more distal radii, with no significant differences observed between any groups beyond 24 µm (p > 0.05).

**Fig. 5.**
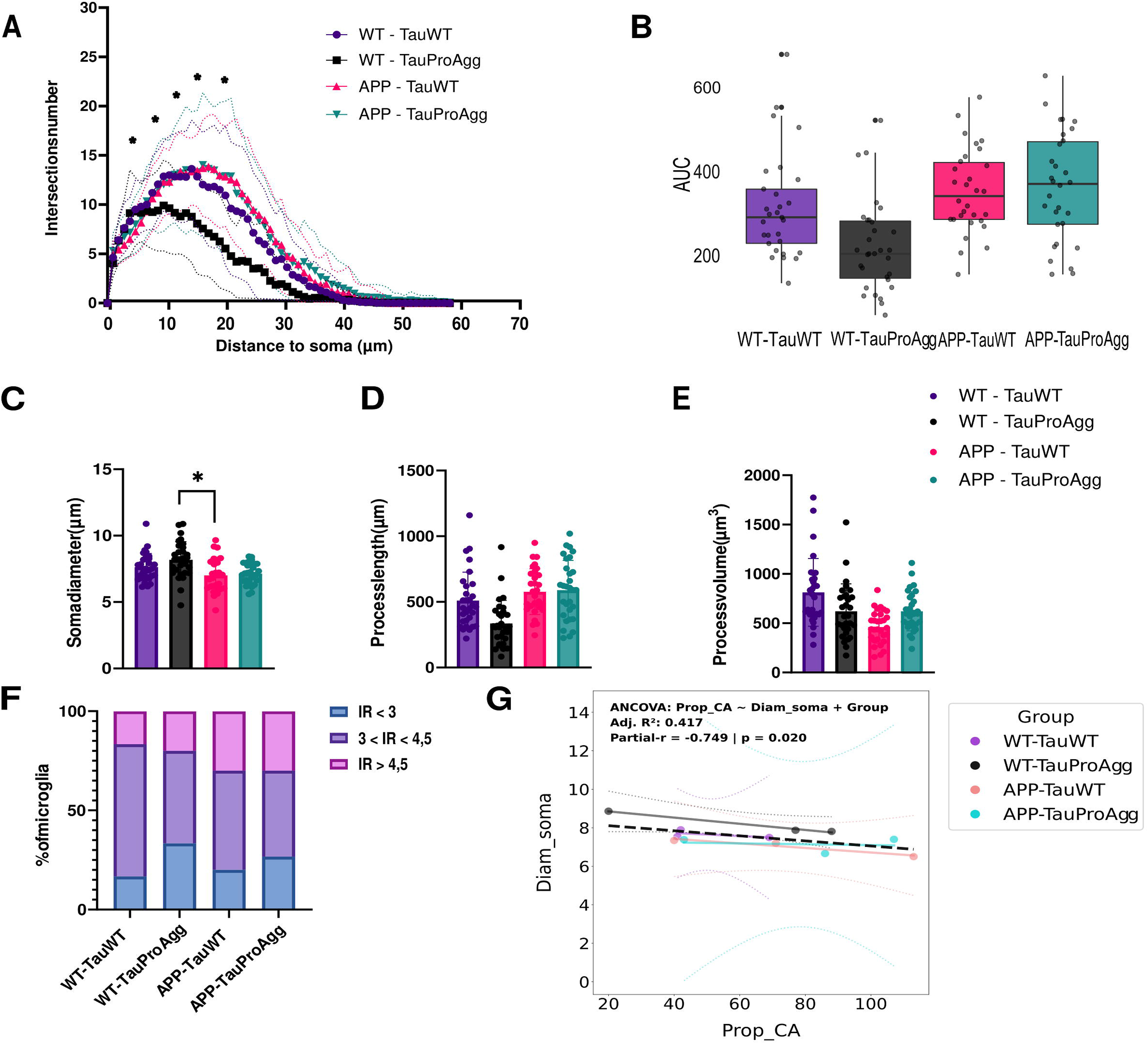
Microglial inflammation in an amyloidosis context. (A) Sholl analysis of microglial morphology near tau spreading site in contralateral DG, quantification of number of intersections vs. distance from soma. Statistical analysis was performed using a linear mixed-effects model (LMM) to account for the hierarchical structure of the data (nested random effects: cells nested within images, nested within animals). The model included Group, Radius, and their interaction as fixed effects. P-values for the fixed effects were estimated using Satterthwaite’s approximation. Post-hoc pairwise comparisons were conducted point-by-point at each radius using Tukey’s HSD adjustment. Asterisks indicate significant differences (p < 0.05)]. Mean ± SEM; n = 10 cells from 3 animals per group. (B-E) Microglial morphological parameters: (B) AUC from Sholl curves, (C) soma diameter (µm), (D) process length (µm), (E) ramification volume (µm³). Mean ± SD; n=30 microglia/group and 3 animals/group; linear mixed-effects model using Satterthwaite’s method, p-values adjusted using Tukey’s post-hoc test for multiple comparisons. (F) Repartition of microglia in function of the IR (IR=Sholl peak / number of primary intersections). Chi2 test. (G) Correlations between tau spreading in CA microglial soma diameter. Data were analyzed using ANCOVA with a common slope model (non-significant interaction removed; p=0.736). Partial R values indicate the correlation between Microglial soma diameter and CA_Propagation across all groups (*p < 0.05).

Sholl AUC did not differ significantly between groups, although values in Tau^ProAgg^-injected WT mice tended to be lower than in the other groups (Fig. 5B; linear mixed-effects model with Satterthwaite’s method, [F(3,9.45)=1.96, p=0.188]). The distribution of microglia across ramification classes was also unchanged (Fig. 5F; chi-square, X2=5.664, df=6, p=0.4619).

Among the morphometric parameters, only soma diameter differed significantly: it was smaller in Tau^WT^-injected APP mice than in Tau^ProAgg^-injected WT mice (Fig. 5C; [F(3,8.01)=4.33, *p<0.05]; Tukey post-hoc, [t(8.08)=-3.24, *p<0.05]). Process length (Fig. 5D; [F(3,8)=2.5, p=0.134]) and process volume (Fig. 5E; [F(3,8)=1.98, p=0.195]) did not differ significantly between groups. Finally, tau spreading to the CA region correlated negatively with microglial soma diameter (Fig. 5G; ANCOVA, Partial-r=-0.749, *p<0.05; the interaction term was not significant and was therefore excluded [p=0.733]).

## DISCUSSION

Cognitive symptoms found in AD are highly correlated with the progression of tau pathology in the brain.^2^ Tau pathology is known to follow a stereotypical anatomical trajectory, initially affecting the entorhinal cortex, then progressing to the hippocampus and eventually invading widespread neocortical regions. Numerous studies have sought to identify factors influencing this propagation, with the ultimate goal of discovering targets that could delay or even halt the spread of tauopathy.

In this study, we developed a novel *in vivo* model relying on the injection of AAVs to overexpress human tau protein in the DG - either in soluble or aggregation-prone form - and on the assessment of tau spreading in the contralateral DG. Using AAV constructs including a GFP reporter gene, *in situ* hybridization and a V5-tagged version of tau to unequivocally identify transduced-cells as well as transgenic tau, we established the relevance of our experimental paradigm to study trans-synaptic tau spreading (Fig. 1).

We then used AAVs expressing three tau variants: wild-type human tau (Tau^WT^), tau carrying the P301L mutation (Tau^P301L^), and an aggregation-prone variant co-expressing wild-type tau with a deltaK280 repeat-domain fragment (Tau^ProAgg^). Mice injected with the aggregation-prone form of tau showed greater tau spreading in both the contralateral DG and CA1-3 regions than those injected with Tau^WT^. Tau aggregation was confirmed in the ipsilateral hippocampus using AT100 immunostaining, whereas little to no aggregation was observed contralaterally. These findings are compatible with the notion that seed-competent tau species, rather than mature fibrillar aggregates, mediate trans-synaptic propagation, an interpretation that aligns with the prion-like hypothesis proposed for misfolded proteins such as PrPSc.^45^ This view has recently received direct structural support: tau filaments extracted from the brains of individuals with Alzheimer’s disease or corticobasal degeneration, injected into wild-type mice, seed the assembly of mouse tau filaments that adopt the same atomic structure as the seeds, establishing that distinct tau folds behave as prion strains.^46^ We note, however, that AT100 immunoreactivity did not differ significantly between Tau^P301L^ and Tau^ProAgg^, so aggregation as measured by AT100 does not by itself account for the difference in spreading between these two constructs. Because distinct tau folds differ in the efficiency with which they are taken up, templated and released^46^, the two constructs may generate assemblies that AT100 detects comparably but that propagate with different efficiency. Testing this would require structural characterization of the assemblies formed by each construct, which cryo-electron microscopy now makes feasible.

Contrasting results were recently reported by Rodrigues et al.,^47^ who used transgenic mice expressing either pro-aggregation or anti-aggregation forms of tau in the entorhinal cortex. They found that neither variant induced spontaneous spreading of tau unless exogenous tau was introduced via AAV transduction, and even then, no differences in spreading were observed between the two tau variants. The authors concluded that tau spreading is independent of its propensity to form aggregates. This discrepancy with our study may be explained by key differences in experimental design. With little aggregation present in the brain, their model likely reflects the early stages of AD pathology, whereas our model - relying on overexpression - may better mimic later stages of the disease. Considering the hypothesis that microglia can engulf and partially degrade tau, generating seed-competent species,^12,36^ it is plausible that the level of expression and the time of exposure are two crucial factors in the accumulation of sufficient tau seeds to allow propagation. It is worth noting that Dujardin et al. recently showed that tau spreading varies depending on the mouse strain used.^48^ In light of this study, it appears that the C57Bl6J background on which we performed our *in vivo* experiments is not the most permissive one for tau spreading, therefore adapting our model to more susceptible strains could provide additional insights.

Given the growing body of literature implicating microglia in tau propagation, we analyzed microglial morphology using Sholl analysis in the contralateral DG at the level where synaptic inputs from transduced mossy cells converge. In all groups, including GFP controls, microglia displayed a ramified morphology without overt signs of activation (Fig. 3), and no significant difference was detected between groups. This analysis was restricted to a small region containing the synapses between mossy cells projecting from the ipsilateral DG and the contralateral post-synaptic neurons of the polymorphic layer, and no uninjected control group was included. Extending the analysis to the whole contralateral hippocampus and comparing it with uninjected WT mice would be needed before drawing conclusions about the microglial response in this model.

The enhanced spreading observed in Tau^ProAgg^-injected mice prompted us to investigate whether extracellular vesicles (EVs) could be involved. We isolated EVs from hippocampal tissue following injection of Tau^WT^ and Tau^ProAgg^ AAVs and found no difference between groups in EV morphology, size distribution or number (Suppl. Fig. 1B-C). Moreover, AT8-positive EVs were significantly more numerous in preparations from Tau^ProAgg^-injected mice. This is consistent with a contribution of EVs to the dissemination of pathological tau. However, the seeding competence of this EV-associated tau was not assessed, and the EV preparations were obtained from pooled, bilaterally injected tissue at a different timepoint from the spreading experiments. These findings are in line with other studies linking tau spreading, microglial activation and EVs.^12,36,49,50^ Determining whether these EVs originate from microglia or from another cell type is an important question that will require further investigation.

To further explore the interaction between tau spreading and amyloidosis, we applied our experimental model to APP/PS1dE9 mice. Due to the high mortality rate in this strain, especially around the age of 4 months, we modified our protocol by injecting 5-month-old mice with a reduced AAV dose and analyzing spreading one month after injection. Under these conditions, no statistically significant differences in tau spreading were observed between groups. This was unexpected, given prior studies demonstrating enhanced tau propagation in the presence of amyloid pathology. However, in this experiment, only a small number of amyloid plaques were detectable, consistent with the relatively young age of the animals. Furthermore, tau aggregation levels were ∼2-fold lower in this experiment than in the first one (Fig. 2 and 4, AT100 positive area/µm2 respectively between 0.1 and 0.2 and between 0.05 and 0.1), most probably because of the reduced viral dose and the shorter post-injection interval. This may explain the absence of an amyloid-related effect on tau spreading. A similar interpretation applies to the neurodegeneration results: while our first experiment (three months post-injection) revealed significant cell loss, no such effects were seen at one month, underscoring the importance of experimental timing.

Although no group-level difference in spreading was detected at one month, the correlations observed in this cohort suggest a possible axis linking the two pathologies. Tau aggregation in the ipsilateral hippocampus was about twice as high in APP mice as in their WT littermates (Fig. 4E), amyloid plaque load correlated with ipsilateral tau aggregation in Tau^ProAgg^-injected APP mice (Fig. 4J, L), and ipsilateral tau aggregation correlated with spreading to the contralateral CA region (Fig. 4G). Taken together, these observations are compatible with the hypothesis that amyloid deposition promotes tau aggregation at the injection site and that this, in turn, favours trans-synaptic spreading. If so, amyloidosis would be expected to accelerate the kinetics of propagation rather than to change its endpoint, and a difference between APP and WT mice should emerge at later time points, as plaque burden increases. Two observations temper this interpretation. First, the same relationship was not found for spreading to the DG (Fig. 4K), which is the primary readout of our model. Second, these correlations are exploratory, were obtained in small groups and were not corrected for multiple comparisons; they generate a hypothesis rather than test one. A longitudinal design sampling several post-injection intervals in the same cohorts would be required to establish whether amyloidosis accelerates tau propagation.

Finally, microglial morphology analysis in APP mice revealed that WT littermates injected with Tau^ProAgg^ AAVs seemed to result in microglial activation, with larger soma and reduced process length, area, and volume as well as diminished arborization complexity (as shown by Sholl analysis, Fig. 5A-E). The lack of difference between APP mice injected with either Tau^WT^ or Tau^ProAgg^ AAVs suggests that Aβ burden induces microglial changes - resembling hyper-ramification - prior to AAV injection, and that tau overexpression does not further affect this hyper-ramified morphology.

## LIMITATIONS

Several limitations should be considered. First, EV cargo was assessed solely with the phospho-specific antibody AT8, so our analysis captures hyperphosphorylated tau only; whether other species - non-phosphorylated, truncated or oligomeric tau - are also loaded into EVs, and in what proportion, remains to be determined. Second, EVs were isolated from pooled, bilaterally injected tissue at a different timepoint from the spreading experiments, and their cellular origin was not determined. Third, spreading was assessed at a single post-injection interval in each experiment - three months in C57BL/6J mice and one month in APP/PS1dE9 mice - so our data provide a snapshot rather than the kinetics of propagation. How the differences between tau variants, and between APP and WT mice, evolve over time, and how they relate to the progression of tau-induced neurodegeneration, cannot be addressed by this design; a longitudinal study sampling several post-injection intervals would be required. Finally, given the marked ipsilateral atrophy observed in Tau^ProAgg^ mice, a contribution of passive tau release from degenerating neurons to the contralateral signal cannot be excluded.

## CONCLUSION

In summary, we established a robust and spatially defined *in vivo* model to study trans-synaptic tau spreading. Our results indicate that tau aggregation enhances spreading, and that aggregation-prone tau is accompanied by a marked enrichment of phosphorylated tau in brain-derived extracellular vesicles, identifying these vesicles as a plausible vehicle for its dissemination. We also show that tau overexpression alone alters microglial morphology, although such effects appear to be diminished or masked in the presence of amyloidosis. These findings highlight the complex interplay between tau pathology, microglia, and amyloid burden, and provide a platform for future studies aiming to dissect the mechanisms of tau propagation and identify therapeutic targets.

## Supporting information

Spplemental Materials and Methods

## LIST OF ABBREVIATIONS

AAV: Adeno-Associated Virus
AD: Alzheimer’s Disease
Aβ: Amyloid-beta peptide
APP: Amyloid Precursor Protein
AUC: Area Under the Curve
bGHpA: Bovine Growth Hormone polyadenylation signal
CA: Cornu Ammonis
DG: Dentate Gyrus
Evs: Extracellular Vesicles
GFP: Green Fluorescent Protein
HSPGs: Heparan Sulfate Proteoglycans
ISH: In Situ Hybridization
LMM: Linear Mixed-effects Model
LRP1: Low-Density Lipoprotein receptor-related Protein 1
ML: Molecular Layer
PL: Polymorphic Layer
PFA: Paraformaldehyde
PS1: Presenilin 1
ROI: Region Of Interest
SYN2: Synapsin 2 promoter
V5: V5 tag
WT: Wild-Type
ANCOVA: Analysis of Covariance
EM: Electron Microscopy
IR: Ramification Index
SEM: Standard Error of the Mean

## STATEMENTS

## Ethics approval

All procedures involving animals were performed following the French regulation (EU directive 86/609 – French Act Rural Code R214–87 to 131). The animal facility was approved by veterinarian inspections (authorization n◦B92-032-02) and complied with Standards for Humane Care and Use of Laboratory Animals of the Office of Laboratory Animal Welfare (OLAW – n◦A5826-01). All procedures received approval from the local ethical committee (Comité d’Ethique en Expérimentation Animale CEtEA) and the French Ministry of Research (APAFIS#<u>612-2015050514376362</u>).

## Availability of data and materials

All data generated or analysed during this study are included in this published article and its supplementary information files.

## Authors’ contributions

PL, CD and A-PB designed the experiments; PL, CD, CA, M-CG, GA, MG, SB, DF, MK, KC and A-PB performed the experiments; ND, FP, CJ, ASH contributed to data acquisition, data processing and results interpretation; PL, CA, MK, KC, A-PB interpreted the results; PL and A-PB wrote the manuscript with input from CD, CA, A-SH, MK, GB and KC. A-PB conceived the project, obtained funding and provided supervision. All authors reviewed the final version of the manuscript.

## Acknowledgements

We acknowledge the DRCM Microscopy Facility-MiP (Université Paris Cité & Université Paris-Saclay, Inserm, DRCM/IBFJ CEA, Stabilité Génétique Cellules Souches et Radiations, F-92260 Fontenay-aux-Roses, France) for the acquisition of high-resolution images using the confocal spinning disk microscopy.

