## Supplemental Figure 1 for "An AAV model of tauopathy targeting the mouse dentate gyrus reveals greater trans-synaptic tau spreading with aggregation-prone tau variants"

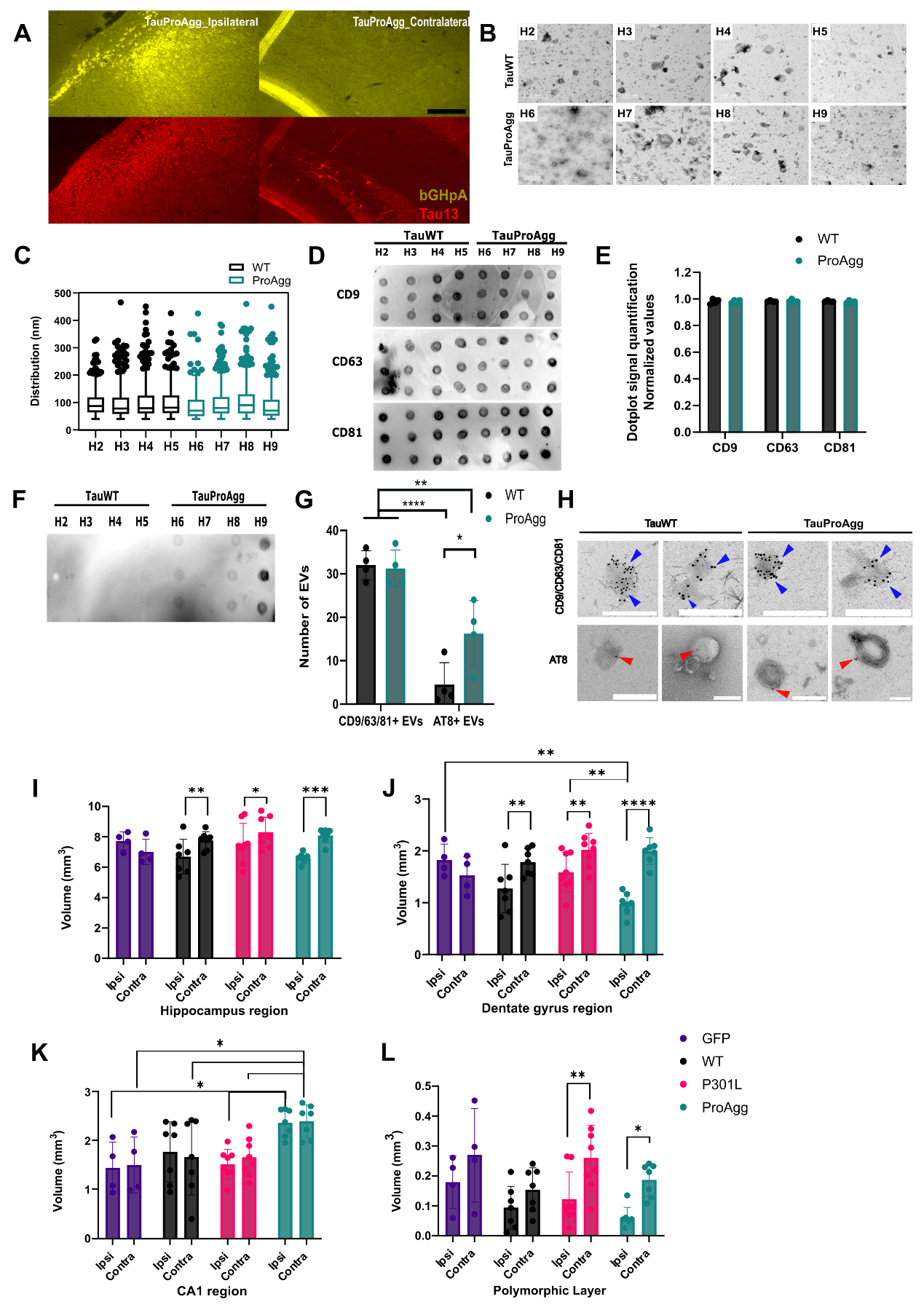

Suppl. Fig.1 Extracellular vesicles characterization and neurodegeneration in DG model.

**(A)** *In situ* hybridization using bGH polyA probe (yellow) specific for the viral genome combined with Tau13 immunofluorescence (red). Scale bar = 50µm. **(B)** Electron micrographs of EVs from each sample. **(C)** EV size distribution of each sample in mice injected with Tau^WT^ (black) or with Tau^ProAgg^ (cyan); each dot representing one EV. **(D)** Dot blot (left panel) with CD9 (top), CD63 (middle) and CD81 (bottom) antibodies on Tau^WT^ (left) and Tau^ProAgg^ (right) samples, and **(E)** dot blot signal quantification (right panel); Mean ± SD; n=4/group; Two-way ANOVA with Sídák's multiple-comparisons test. Average number of EVs obtained in each case plotted with each dot representing one mouse. **(F)** Dot blot with AT8 antibody on Tau^WT^ (left) and Tau^ProAgg^ (right) samples. **(G)** Quantification of CD9/63/81- and AT8-positive EVs. Each dot represents one mouse. Mean ± SD; n=4/group; Two-way ANOVA with Sídák's multiple-comparisons test; * p<0.05, ** p< 0.01, **** p < 0.001. **(H)**Electron micrographs of EVs immunogold-labeled for CD9/CD63/CD81 (top panels) or AT8 (bottom panels) from Tau^WT^- (left) or Tau^ProAgg^ hippocampus (right). Cyan arrows indicate CD9/CD63/CD81 labeling; red arrows indicate AT8 staining.**(I-L)** Volume extrapolated from surface measurement on a sample of section of both ipsilateral and contralateral side for (I) hippocampus, (J) DG, (K) CA1 region, and (L) polymorphic layer of the DG. Data are presented as mean ± SD, n=8/group; Two factors ANOVA with Sídák's multiple comparisons test; *p< 0.05, **p< 0.01, ***p <0.001, ****p<0.0001.
