## Supplemental Figure 2 for "An AAV model of tauopathy targeting the mouse dentate gyrus reveals greater trans-synaptic tau spreading with aggregation-prone tau variants"

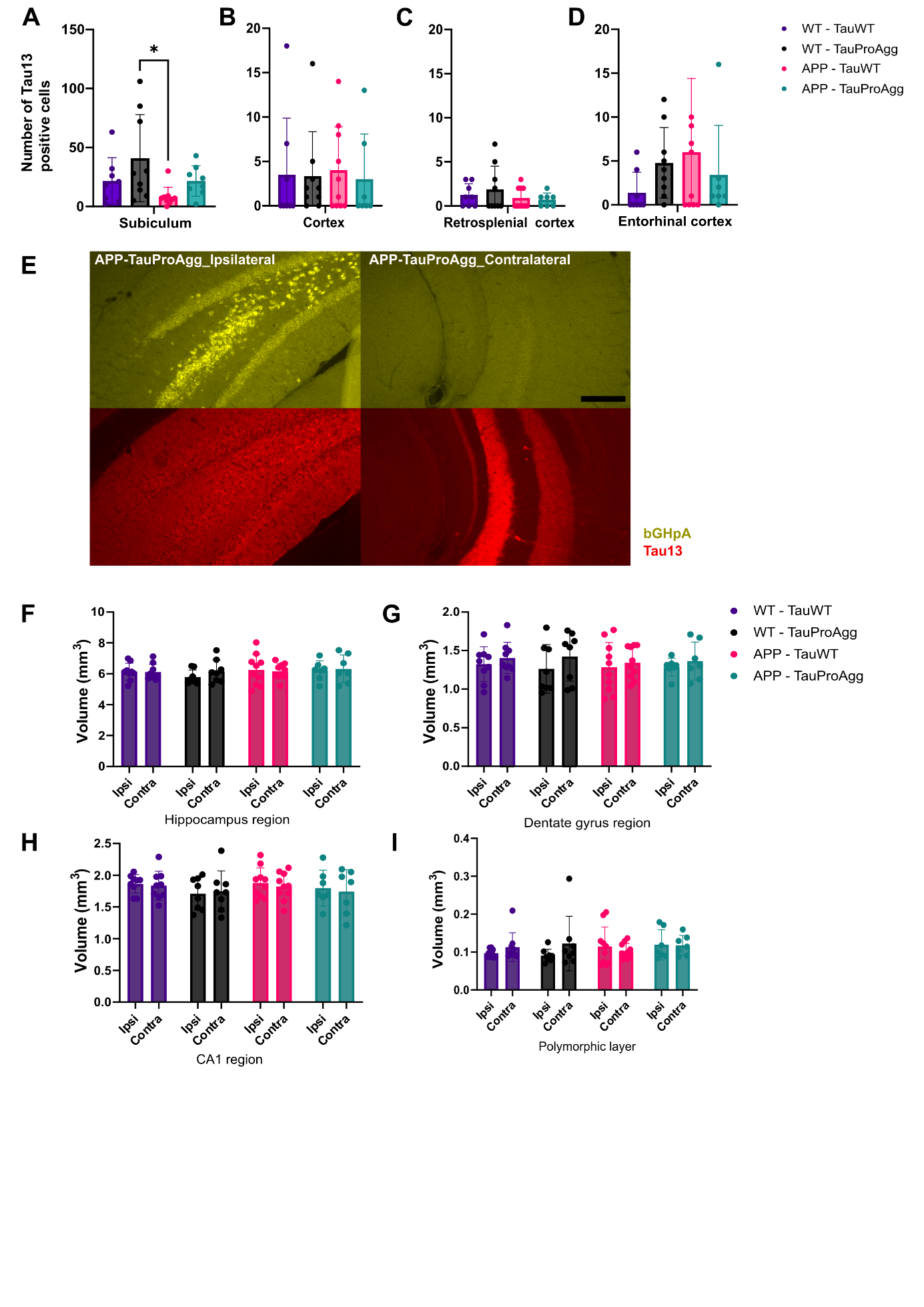

Suppl. Fig.2 Tau spreading in additional brain regions and characterization of the DG model in APP mice.

**(A-D)** Quantification of Tau13-positive neurons in contralateral subiculum, cortex, retrosplenial cortex (Rsple), and entorhinal cortex (Ent) in WT mice injected with Tau^WT^ (purple), WT mice injected with Tau^ProAgg^ (black), APP mice injected with Tau^WT^ (pink) and APP mice injected with Tau^ProAgg^ (cyan). Mean ± SD; n=8/group. Kruskall-Wallis test with Dunn’s multiple-comparisons test. **(E)** In situ hybridization with bGH polyA probe (yellow) combined with Tau13 immunofluorescence (red). Scale bar = 50µm. **(F-I)** Volumetric analysis (mm3) of ipsilateral and contralateral (F) hippocampus, (G) DG, (H) CA1 region, (I) polymorphic layer of the DG. Mean ± SD; n=8/group; Two-way ANOVA with Sídák's multiple-comparisons test.
