## Supplementary material for "An AAV model of tauopathy targeting the mouse dentate gyrus reveals greater trans-synaptic tau spreading with aggregation-prone tau variants": Spplemental Materials and Methods

**SUPPLEMENTARY FIGURES AND TABLE**

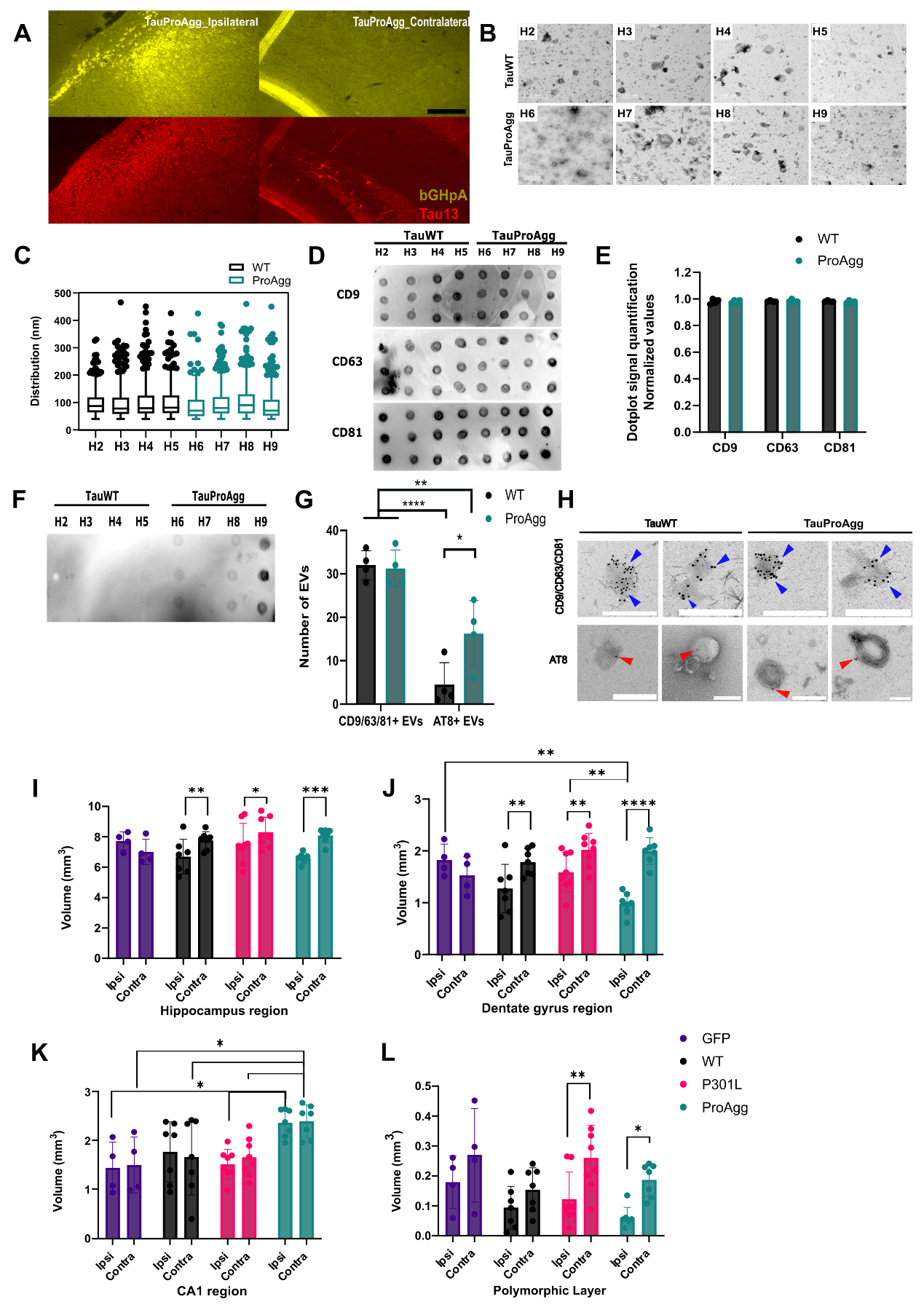

Suppl. Fig.1 Extracellular vesicles characterization and neurodegeneration in DG model.

**(A)** *In situ* hybridization using bGH polyA probe (yellow) specific for the viral genome combined with Tau13 immunofluorescence (red). Scale bar = 50µm. **(B)** Electron micrographs of EVs from each sample. **(C)** EV size distribution of each sample in mice injected with Tau^WT^ (black) or with Tau^ProAgg^ (cyan); each dot representing one EV. **(D)** Dot blot (left panel) with CD9 (top), CD63 (middle) and CD81 (bottom) antibodies on Tau^WT^ (left) and Tau^ProAgg^ (right) samples, and **(E)** dot blot signal quantification (right panel); Mean ± SD; n=4/group; Two-way ANOVA with Sídák's multiple-comparisons test. Average number of EVs obtained in each case plotted with each dot representing one mouse. **(F)** Dot blot with AT8 antibody on Tau^WT^ (left) and Tau^ProAgg^ (right) samples. **(G)** Quantification of CD9/63/81- and AT8-positive EVs. Each dot represents one mouse. Mean ± SD; n=4/group; Two-way ANOVA with Sídák's multiple-comparisons test; * p<0.05, ** p< 0.01, **** p < 0.001. **(H)**Electron micrographs of EVs immunogold-labeled for CD9/CD63/CD81 (top panels) or AT8 (bottom panels) from Tau^WT^- (left) or Tau^ProAgg^ hippocampus (right). Cyan arrows indicate CD9/CD63/CD81 labeling; red arrows indicate AT8 staining.**(I-L)** Volume extrapolated from surface measurement on a sample of section of both ipsilateral and contralateral side for (I) hippocampus, (J) DG, (K) CA1 region, and (L) polymorphic layer of the DG. Data are presented as mean ± SD, n=8/group; Two factors ANOVA with Sídák's multiple comparisons test; *p< 0.05, **p< 0.01, ***p <0.001, ****p<0.0001.

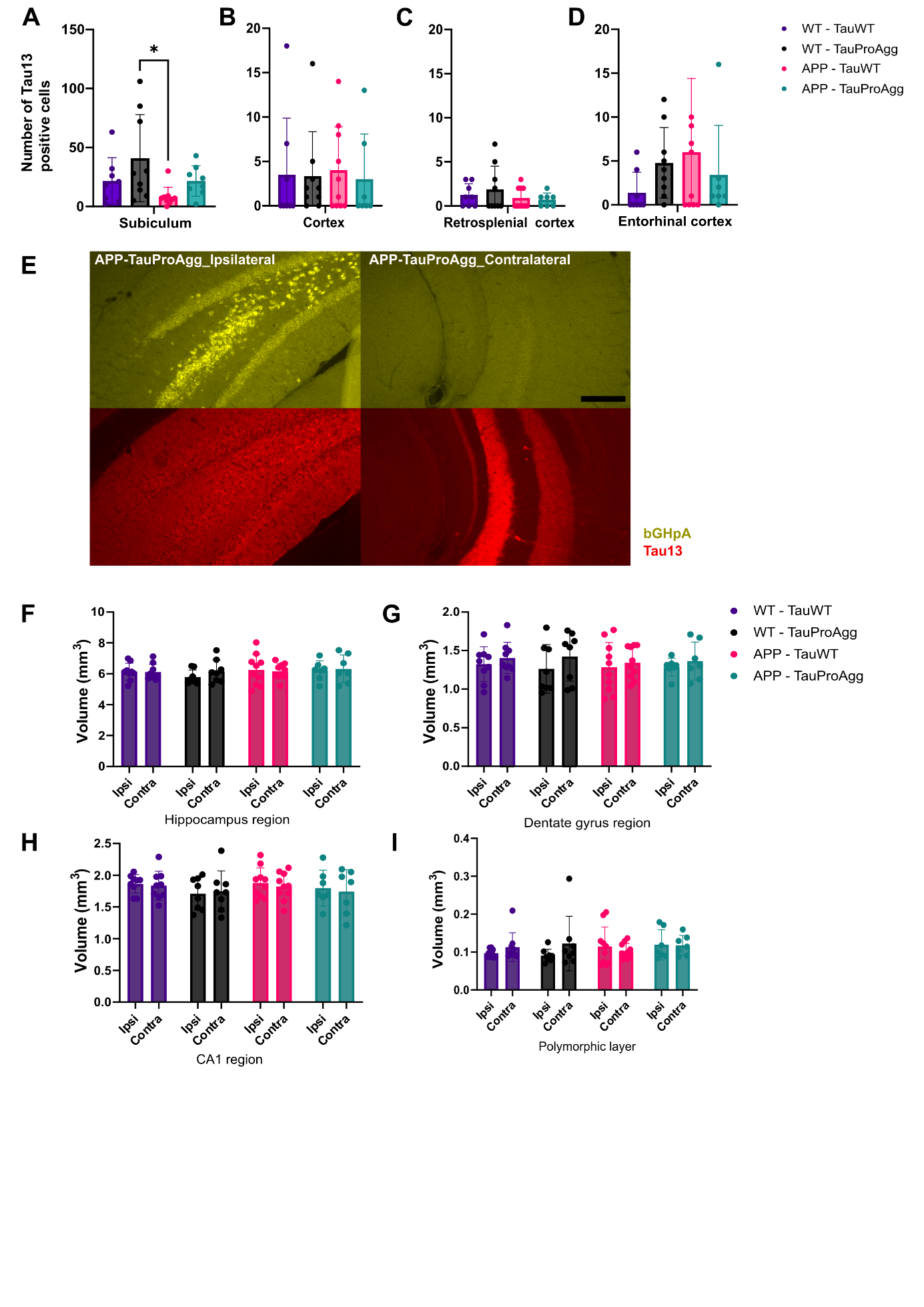

Suppl. Fig.2 Tau spreading in additional brain regions and characterization of the DG model in APP mice.

**(A-D)** Quantification of Tau13-positive neurons in contralateral subiculum, cortex, retrosplenial cortex (Rsple), and entorhinal cortex (Ent) in WT mice injected with Tau^WT^ (purple), WT mice injected with Tau^ProAgg^ (black), APP mice injected with Tau^WT^ (pink) and APP mice injected with Tau^ProAgg^ (cyan). Mean ± SD; n=8/group. Kruskall-Wallis test with Dunn’s multiple-comparisons test. **(E)** In situ hybridization with bGH polyA probe (yellow) combined with Tau13 immunofluorescence (red). Scale bar = 50µm. **(F-I)** Volumetric analysis (mm3) of ipsilateral and contralateral (F) hippocampus, (G) DG, (H) CA1 region, (I) polymorphic layer of the DG. Mean ± SD; n=8/group; Two-way ANOVA with Sídák's multiple-comparisons test.

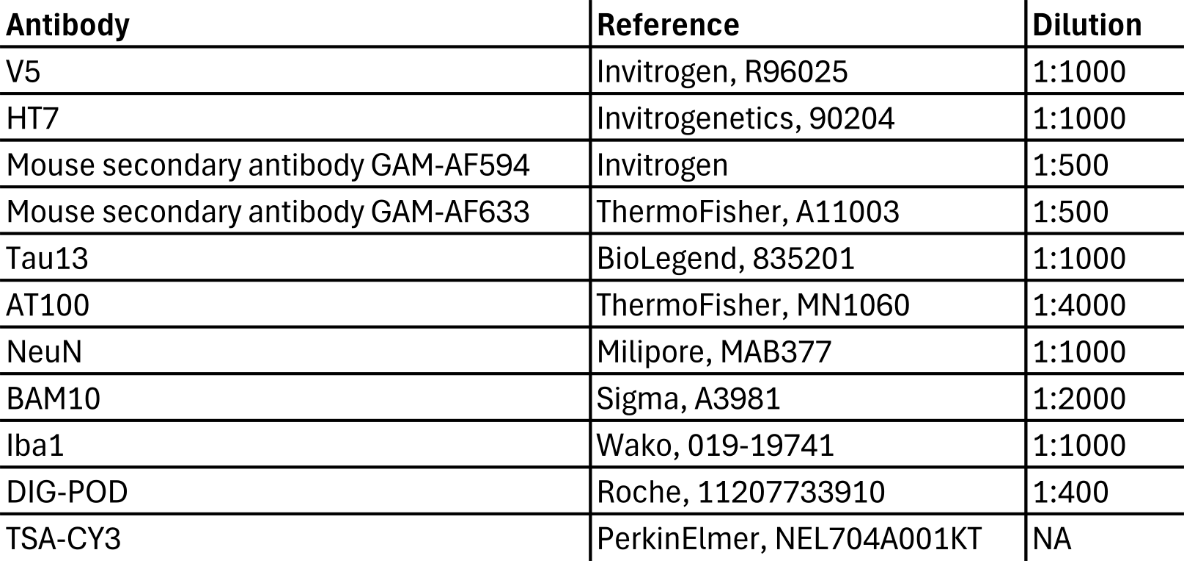

Suppl.Table1. Histological product references and dilutions

**SUPPLEMENTARY MATERIALS & METHODS**

*Viral vectors design and production*

AAV9 particles were produced by transient transfection of HEK-293T cells with a three-plasmids system as previously described.^1^ Three days after transfection, viral particles were harvested in cell lysate and supernatant, purified by ultracentrifugation on a Iodixanol discontinuous density gradient, and subsequently desalted and concentrated by tangential flow ultrafiltration. AAV titers were estimated by QPCR using a plasmid of known concentration as standard and expressed as genome content per mL (GC/mL). The resulting AAV vector batches were aliquoted and stored at -80°C until further use.

*AAV stereotaxic injections*

Mice were premedicated with buprenorphine (0.075mg/kg, Vetergesic^®^) and anesthetized with an intraperitoneal injection of a mix of ketamine (100mg/kg, Imalgène 1000^®^) and medetomidine (0.25mg/kg, Domitor^®^). Local analgesia was obtained by subcateneous injection of lidocaine (5mg/kg, Lurocain^®^) 5 minutes before scalp incision and drilling a hole using a milling cutter at the injection site. After injection in 10minutes of AAV diluted in a total volume of 2µL, the injection canula was left in place for an additional 5 minutes to allow sufficient diffusion of the solution. The injection canula was finally slowly withdrawn, the skin disinfected and stapled. Following stereotaxic surgery, reversion of anesthesia was induced by subcutaneous injection of atipamezole (0.25mg/kg, Antisedan^®^), the animals were placed in a thermostatically controlled recovery cage and monitored until complete recovery. The injection canula was then slowly withdrawn, the skin disinfected and stapled. Mice were rehydrated with a subcutaneous injection of normal saline and kept under constant monitoring in a thermo-regulated box until complete recovery from anesthesia.

*In Situ Hybridization and immunolabeling*

Brain slices were rinsed in PBS1X solution and then saturation of non-specific binding sites was performed by incubation at 68°C in a pre-hybridization buffer solution composed of SSC5X (0.75M NaCl, 0.075 sodium citrate pH7) containing 50% of deionized formamide. The hybridization probe was diluted 1:100 in a hybridization buffer containing 50% formamide, 5X SSC, 0.2mg/mL yeast tRNA, 0.1% Tween20, and 1mg/mL Denhardt’s solution in distilled water. The hybridization buffer containing the probe was denatured at 80°C for 5 minutes and stored immediately on ice to avoid renaturation. Brain slices were then incubated overnight at 70°C in the hybridization buffer. Several washings were then performed with decreasing concentrations of SSC (from 2X to 0.2X) and formamide (from 50% to 0%) and decreasing temperatures (from 70°C to 60°C). Brain slices were then rinsed in TNT buffer (0.015M of NaCl, 0,1M of pH7.5 Tris-HCl, 0,1% Tween20) at room temperature and subsequently incubated for 1 hour at room temperature in a blocking solution containing 0.5% Blocking Reagent (Roche Diagnostics GmbH, Mannheim, Germany) and 2% normal goat serum in TNT buffer. Brain slices were then incubated overnight at 4°C in the blocking solution containing 1:400 anti-digoxygenin antibody, rinsed three times in TNT buffer, incubated with TSA-CY3 (PerkinElmer, Villebon-sur-Yvette, France) to reveal the probe labeling, rinsed three times in TNT buffer and three times in PBS1X and finally fixed with 4% PFA for 5 minutes. Immunostaining against V5 or Tau13 was then performed as described in the main materials & methods section.

*Image analysis*

To measure tau aggregation, we quantified the AT100-positive area using ImageJ software on each brain section for each animal after defining regions of interest (ROI) corresponding to ipsilateral and contralateral hippocampus. To determine AT100-positive area we applied a threshold defined on control slices. Shrinkage of different sub-regions of the hippocampus was used as a proxy to evaluate neurodegeneration: we quantified the area of different ROIs (whole hippocampus, CA1 layer, DG, polymorphic layer of the DG) on all the brain slices from –2.5mm to +3.5mm relative to bregma for each animal on one series of sections. Then we extrapolated the volume of these regions thanks to the following formula.

$$\boldsymbol{volume=}\sum\boldsymbol{ROI area x thickness of slices x number of series}$$

Finally, to evaluate the density of amyloid plaques, we developed a macro algorithm on ImageJ using a threshold to define the specific labeling of BAM10 allowing us to measure the size and the number of the plaques. These parameters were measured in the DG and hippocampus both on ipsilateral and contralateral sides on all the brain slices from –2.5mm to +3.5mm from the bregma for each animal on one series of sections labelled with BAM10 antibody.

*Isolation and characterization of brain-derived extracellular vesicles*

Mice were injected with buprenorphine (0.3 mg/kg) 30 minutes before euthanasia with a lethal dose of ketamine (220 mg/kg) and medetomidine (0.7 mg/kg), then brain were harvested and dissected on ice in order to collect hippocampi and cortical tissues separately. Hippocampal (average tissue weight: 73±4 mg) and cortical (average tissue weight: 169±10 mg) tissues were resuspended in 150 µL or 300 µL, respectively, of dissociation buffer (10 mM HEPES-OH pH 7.4, 137 mM NaCl, 5 mM KCl, 1 mM MgCl_2_, 1.8 mM CaCl_2_) containing 2 mg/mL Collagenase D and 40 U/mL DNase I, and incubated for 10 min at 37°C with low-speed (300 rpm) agitation on a thermomixer (Eppendorf®). The reaction mixes were immediately put on ice and an equal volume of ice-cold dissociation buffer containing protease (1 mM PMSF, Complete Roche®) and phosphatase (PHOS-stop, Roche®) inhibitors were added to stop the reaction. Tissues were very gently dissociated on ice by slow repeated pipetting using a trimmed tip. Tissues and cell debris were removed by centrifugation at 500 x g for 5 min and the supernatant further cleared by centrifugation at 2,000 x g for 10 min and by passage through a syringe filter (0.45 µm). The filtrates containing EVs were loaded on qEVsingle (gen2) 35nm size exclusion chromatography columns (Izon, Lyon France) and processed using Izon automatic fraction collector (AFC V2) and following the manufacturer’s recommendations. EVs were eluted using PBS and recovered in fractions 2 and 3 (150 µL each) which were pooled, aliquoted and stored at -80°C until use. EVs preparations were characterized by negative-staining electron microscopy (EM), as described before ^2^ with minor modifications. EVs were mixed with an equal volume of 4% PFA and incubated for 20 min at 4°C. Fixed EVs were then applied to formvar carbon-coated EM grids (EMS, Hatfield, PA, USA) and allowed to adsorb for 20 min at RT. EM grids were washed by sequential transfers on drops of PBS and then on drops of water. Following negative-staining with 1% Uranyl acetate or Uranyless EM-stain (Delta Microscopies, Mauressac, France) for 5 min at RT, samples were imaged in a Jeol 1400 transmission electron microscope (Jeol, Croissy-sur-Seine, France). Images were recorded with a Gatan Orius CCD camera (Gatan, Pleasanton, CA) and processed with ImageJ software. For immunogold-EM analysis, EVs were fixed and adsorbed on EM grids as described above. Residual aldehyde groups were inactivated for 10 min at RT with 50mM glycine in PBS, followed by washes on drops of PBS. Grids were blocked with 5% BSA in PBS for 30min at RT and subsequently rinsed on several drops of immunoblocking buffer (PBS containing 0.1% BSA-c, Aurion, Wageningen, The Netherlands). EVs were then permeabilized with an immunoblocking buffer containing 0.02% of Triton-X100 for 10 min at RT, followed by washes in several drops of immunoblocking buffer. EM grids were then incubated for 1h at RT either with a mix of rabbit recombinant anti-CD9, CD63, CD81 antibodies (Abcam Ltd, Cambridge, UK), or with a recombinant AT8 antibody (a generous gift from Dr Luc Bousset). All primary antibodies were diluted at 1/40 in immunoblocking buffer. EM grids were then washed on several drops of immunoblocking buffer and incubated for 1h at RT with anti-rabbit secondary antibody conjugated to 6nm gold particles (Aurion, Wageningen, The Netherlands) diluted at 1/40. EM grids were washed in several drops of immunoblocking buffer and PBS and then incubated for 5min in PBS containing 2.5% glutaraldehyde for 5min at RT. EM grids were then washed in several drops of PBS and water before negative staining as described above with Uranyless EM-stain (Delta Microscopies). Semi-quantitative analysis of the immunogold labelling was achieved by analyzing each EM grid under the electron microscope for 20 to 30 min, making sure to cover and image different areas of the grid. CD9/63/81 and AT8 positives-EVs were identified and counted manually during the imaging session and then recounted on the micrographs using ImageJ analysis.

EVs were also analyzed by immunoblotting. To this end, 5 µL of EVs were spotted on nitrocellulose membranes and allowed to dry. The membranes were blocked with 5% BSA in TBST (Tris-Buffered Saline, 0.1% Tween-20) and then incubated overnight at 4°C under low agitation in TBST containing primary antibodies (anti-CD9, CD63, CD81 or AT8) at a 1/5000 dilution. The membranes were then washed in TBST and incubated in TBST containing Horseradish peroxidase (HRP)-conjugated secondary antibodies for 2h at RT under low agitation. After several washes in TBST, membranes were imaged and analyzed using Super Signal West Femto enhanced chemiluminescence (ECL) reagents (Thermo Fisher Scientific, Les Ulis, France) and a Fusion FX7 Imager (Thermo Fisher Scientific).
