## Supplementary figures and images for "An AAV model of tauopathy targeting the mouse dentate gyrus reveals greater trans-synaptic tau spreading with aggregation-prone tau variants"

### Supplemental Table 1

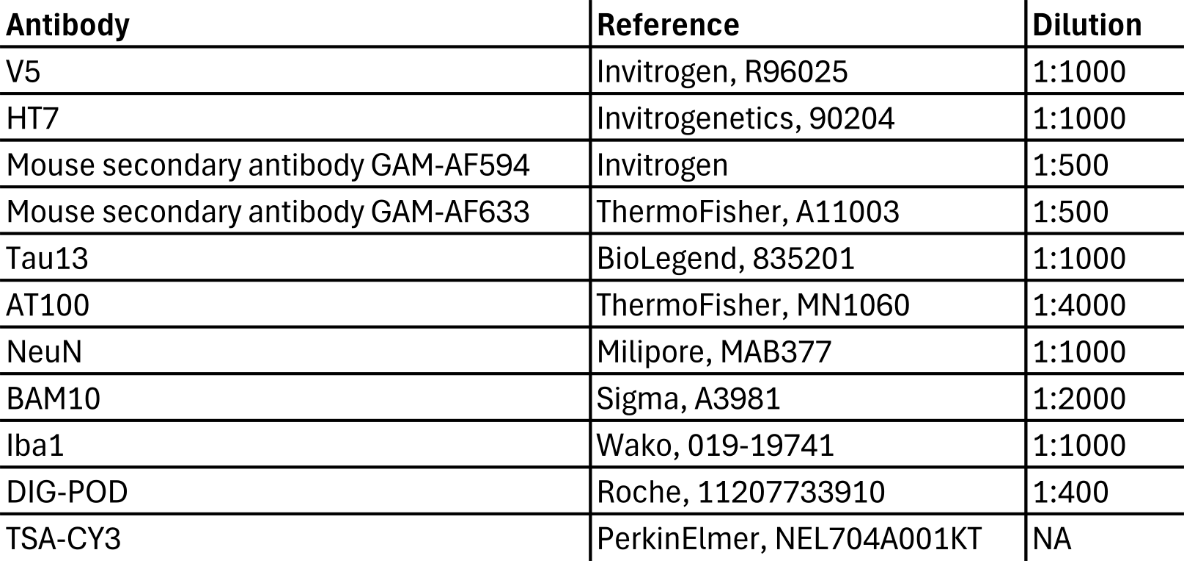

Suppl.Table1. Histological product references and dilutions
